# A concurrent canonical and modified miRNAome pan-cancer study on TCGA and TARGET cohorts leads to an enhanced resolution in cancer

**DOI:** 10.1101/2021.05.18.444694

**Authors:** Rosario Distefano, Luisa Tomasello, Gian Luca Rampioni Vinciguerra, Pierluigi Gasparini, Yujia Xiang, Marina Bagnoli, Gioacchino Paolo Marceca, Paolo Fadda, Alessandro Laganà, Mario Acunzo, Qin Ma, Giovanni Nigita, Carlo M. Croce

## Abstract

MiRNA Epitranscriptomics has placed a new layer of complexity in the cancer field. Despite the fast-growing interest in miRNA editing and shifted miRNA isoforms, a simultaneous study of both modifications in cancer is still missing. Here, we concurrently profiled multiple miRNA modifications, including A-to-I RNA editing and shifted miRNA isoforms, in >13K adult and pediatric tumor samples across 38 distinct cancer cohorts from The Cancer Genome Atlas and The Therapeutically Applicable Research to Generate Effective Treatments datasets. We investigated the differences among canonical miRNAs and the wider miRNAome in terms of expression, clustering, dysregulation, and prognostic standpoint. The combination of canonical miRNAs/miRNA isoforms boosted the quality of clustering results, outlining unique cohorts’ clinical-pathological features. We described modified miRNAs showing opposite dysregulation with respect to their canonical counterparts in cancer, potentially impacting their targetome and function. The abundance of expressed miRNA isoforms directly impacted the activation/deactivation of critical carcinogenesis pathways. Finally, we experimentally validated unique targeting for a shifted and edited miRNA isoform. Our findings outlined once more the importance of going beyond the well-established paradigm of one-mature-miRNA per miRNA arm to elucidate novel mechanisms related to cancer progression.

## INTRODUCTION

MicroRNAs (miRNAs) are a class of small non-coding RNA (ncRNA) molecules of ~21 nucleotides (nts) in length, expressed in eukaryotes, which negatively regulate gene expression at the post-transcriptional level (1). Their dysregulation observed in several diseases (2), including cancer (3), prompted the interest in employing such molecules as diagnostic and prognostic cancer biomarkers (4). Until recently, most studies on miRNAs widely relied upon the miRNA biogenesis paradigm, “one mature miRNA per miRNA precursor arm” (1,5). However, the latest advancements in next-generation sequencing (NGS) technologies unveiled a more complex scenario (6), in which some expressed miRNA molecules differ from their reference sequence (miRBase) (7). These miRNA variants, termed *miRNA isoforms* or *isomiRs*, undergo several RNA modification events, such as RNA Editing (8,9), miRNA sequence alternative cleavage (10,11), and alterations at the DNA level (i.e., Single Nucleotide Polymorphisms – SNPs) (12). The growing interest in such phenomena undermined the before-mentioned well-established paradigm (7).

The Adenosine-to-Inosine (A-to-I) RNA editing (8,9) represents the most abundant RNA editing variant in mammals, affecting splicing and translational machinery. Occurring in double-stranded (ds) RNA regions, it affects both coding and non-coding RNAs (9), including miRNAs (13). RNA editing has been associated with several human diseases (14–16). A single RNA editing event within the miRNA seed region (MSR) could compromise the miRNA-mediated gene regulation process, which in turn may drastically alter the miRNA targetome (13,17). Likewise, *shifted isomiRs*, which may result from imprecise cleavage of the miRNA reference sequence, could induce targetome differentiation (18). Shifted isomiRs may likely result from the imprecise cleavage processing by Drosha, in stark contrast with Dicer, which cleavages at a fixed distance (19). Initially considered as artifacts (20), their function has been recently re-evaluated (21) as they actively interact with mRNAs (10,11,22). Like the A-to-I RNA editing occurring within the MSR, the 5’-end shifting (addition/trimming of nts at 5’-end) may diversify the targetome revealing a more complex role in gene regulation than previously expected (23). Interestingly, the expression profile of shifted isomiRs exhibits high tissue and cell variability (20), with some cases reporting an expression higher than their canonical counterparts (24).

Albeit the rising interest in the miRNA Epitranscriptome, most studies have investigated a subset of miRNA modifications. In the last years, efforts on assessing the biological implications of A-to-I RNA Editing affecting canonical miRNAs have inflated the interest in employing such molecules as potential biomarkers for cancer prognosis and therapy (25,26). Recently, a surge of shifted isomiRs-oriented studies has been emerged (27,28), with the first pan-cancer study (29) profiling 32 tumor cohorts in The Cancer Genome Atlas (TCGA).

Although the studies mentioned above have proven the importance of investigating such miRNA modifications, a concurrent profiling of such miRNA modification events in cancer is still missing. In this work, we relied on miRge 2.0 (30), one of the major pipelines for canonical miRNAs/miRNA isoforms profiling, given its reliability and accuracy in identifying A-to-I RNA editing sites (see Supplementary Information for more details). Here, we concurrently profiled canonical miRNAs, shifted isomiRs, modification events such as A-to-I RNA Editing and SNPs, as well as nucleotide insertions. We processed information at a large scale from the most prominent and reliable cancer public resources, TCGA and The Therapeutically Applicable Research to Generate Effective Treatments (TARGET), analyzing >13K adult and pediatric cancer samples spread across 38 distinct cohorts. In our data, the abundance of expressed annotated isomiRs outmatched by *8*-fold the number of expressed canonical miRNAs (miRBase v22). We observed a predominance of 3’-end shifted molecules among the expressed miRNA modifications, with an expression comparable to the canonical ones. Both miRNA arms (5p and 3p) displayed higher mobility at 3’-end (large trimmings/additions) over the more conservative 5’-end (small trimmings/additions). Interestingly, most predominant SNP forms exclusively characterized the two arms, while the A-to-I RNA editing sites were mainly located within the 1-10 nts region on both arms. At first, we explored the capability of isomiRs to cluster cancer samples across cohorts. We then examined the differences in the abundance of expressed isomiRs across cohorts’ cancer samples, finding several cancer-related pathways enriched significantly. Afterward, we investigated isomiRs from a diagnostic and prognostic standpoint. Finally, we experimentally validated gene targeting for two different isomiRs: a shifted miRNA (with no Single Nucleotide Variants - SNVs) and an edited miRNA (with no sequence shifting). Using a combination of canonical miRNAs and isomiRs, we obtained a higher cluster fragmentation that reflected a more profound clinical-pathological stratification. IsomiRs resulted significantly deregulated across cohorts/cancer tissues, showing a distribution of modification types proportional to the one observed for expressed annotated molecules. Overall Survival (OS) and Relapse Free Survival (RFS) prognostic signatures were significantly enriched with isomiRs over canonical miRNAs, with multiple signatures entirely composed of isomiRs. Finally, we experimentally assessed gene targeting exclusivity by investigating the canonical miR-101-3p and one of its shifted isomiRs in lung adenocarcinoma, and the canonical miR-381-3p and one of its edited form (A-to-I editing at position 4) in breast cancer.

In summary, our findings highlight the importance of considering the broader modified miRNAome, which actively participates in gene regulation and may offer the opportunity to discover novel cancer biomarkers.

## MATERIALS AND METHODS

### Cell lines

HEK-293, A549, H1299, MDA-MB-231, and HCC70 (ATCC) were seeded and grown in RPMI-1641 medium supplemented with 10% of fetal bovine serum and penicillin-streptomycin (100 U/mL penicillin and 0.1 mg/mL streptomycin) (Millipore Sigma). All cell lines were authenticated through the short-tandem repeat profiling method and tested to be free of mycoplasma contamination.

### Cell transfection

HEK-293, A549, H1299, MDA-MB-231, and HCC70 cell lines were plated in a 6- or 12-wells plate 24 hours before transfection. 100 nM of *miR-101-3p* (full label *miR-101-3p mir-101-1_0_0_21M*, see *MiRNA Isoform Nomenclature* for more details) mirVana™ miRNA Mimic, *miR-101-3p (−1|−2)* (full label *miR-101-3p_mir-101-1_−1_−2_20M*, see *MiRNA Isoform Nomenclature* for more details) custom mirVana™ miRNA Mimics, *miR-381-3p* (full label *miR-381-3p_mir-381_0_0_22M*, see *MiRNA Isoform Nomenclature* for more details) mirVana™ miRNA Mimic, and *miR-381-3p_4_A_G* (full label *miR-381-3p_mir-381_0_0_3MG18M*, see *MiRNA Isoform Nomenclature* for more details) custom mirVana™ miRNA Mimics (all by Thermo Fisher Scientific) were transfected using Lipofectamine™ 2000 Transfection Reagent (Thermo Fisher Scientific) diluted in transfection-medium (RPMI-1641 without FBS or antibiotics). mirVana™ miRNA Mimic, Negative Control #1, and Anti-miR™ miRNA Inhibitor Negative Control #1 (both by Thermo Fisher Scientific) were employed as scrambled controls. After 5 hours, transfection-medium was replaced with RPMI-1641 supplemented with 10% fetal bovine serum and penicillin-streptomycin and, only for H1299 cells, with mitomycin C at the concentration of 15μg/ml. After 24 hours or 48 hours, cells were harvested and subjected to Luciferase assay or RNA isolation and protein lysis. See Supplementary Information for more details.

### Validation of canonical and novel miRNA-target by luciferase assay

PsiCHECK-2 vector (Promega) was employed to generate luciferase-based reporters for miRNA-target validation. 3’ UTR of PTGS2, DSC2, UBE2C, and SYT13 genes, containing miRNAs binding sites and ~50-200 nt flanking regions, were amplified by PCR from human genomic DNA (Promega), using the primers listed in Table S1 and then inserted into the psiCHECK-2 vector, downstream to *Renilla* luciferase open reading frame. All inserted sequences were checked via Sanger Sequencing. 500 ng of psiCHECK™-2 vector holding the specific 3’UTR were transfected together with 100 nM of mirVana™ miRNA Mimics or Negative Controls in HEK-293 cells, as described above. After 24 hours, cells were lysed, and *Firefly* (internal control) and *Renilla* enzymatic activity were measured using Dual-Luciferase® Reporter Assay System (Promega) and detected by GloMax® 96 microplate luminometer (Promega), according to the manufacturer’s protocol. The comparison statistical significance was computed using the two-tailed unpaired Student’s t-Test provided by the *t.test* function available in *stats*, an R (v3.4.4) package.

### RNA isolation, reverse transcription, and real-time RT-PCR

The expression of canonical microRNAs, isomiRs, and target genes was analyzed by real-time RT-PCR after designing custom TaqMan® Small RNA Assays for isomiRs detection (Thermo Fisher Scientific). Total RNA was isolated from cells 48 hours post-transfection using TRIzol™ Reagent (Thermo Fisher Scientific) according to the standard protocol and measured with the Nanodrop 2000c instrument (Thermo Fisher Scientific). For specific microRNAs reverse-transcription, cDNA was synthesized from 5 ng of total RNA using TaqMan® Small RNA Assays RT-primers (Thermo Fisher Scientific) with the High-Capacity cDNA Reverse Transcription Kit, 30 min 16 °C, 30 min 42 °C, 5 min 85 °C. MicroRNA real-Time RT-PCR was performed using TaqMan™Fast Advanced Master Mix according to the manufacturer’s protocol, with cataloged and custom TaqMan® Small RNA Assays (Thermo Fisher Scientific): miR-101-3p (assay ID 002253), miR-101-3p (−1|-2) (custom assay), miR-381-3p (assay ID 000571), and miR-381-3p_4_A_G (custom assay). The data were normalized using RNU44 (assay ID 001094).

### Protein lysis and western blotting analysis

Cells were lysed in Lysis buffer (50 mM Tris HCl pH 7.5, 150 mM NaCl, 10% Glycerol, and 0.5% Nonidet P40), supplemented with Protease inhibitors (Millipore Sigma). Then, 15-25 μg of proteins were loaded onto 4-12% Mini-PROTEAN Tris-Tricine Precast Gels or Criterion Tris-Tricine Precast Gels (Bio-Rad), and electro-blotted on nitrocellulose membranes (GE Healthcare Life Science). Later, the membranes were blocked in blocking solution (TBS-0.05% Tween®20/fat-free milk 5% or TBS-0.05% Tween®20/BSA 5%) and incubated overnight at 4°C with all the primary antibodies, anti-Cox2 (ABclonal Technologies and Cell Signaling), anti-DSC2 (ABclonal Technologies), anti-UBE2C (ABclonal Technologies), anti-SYT13 (Thermo Fisher Scientific), and anti-Vinculin (Abcam), previously diluted in fat-free milk 3-5%. The day after, the membranes were washed three times with TBS-0.05% Tween®20 (TBS-T) and incubated with appropriate HRP-conjugated secondary antibodies (Millipore Sigma), one hour at room temperature. After three additional washes, the membranes were assayed with ECL (Millipore Sigma), and the signal was marked and developed on Blue X-ray film (GeneMate) inside a dark room.

### RNA-binding Protein Immunoprecipitation (RIP)

For RNA-binding protein immunoprecipitation (RIP) ~3×10^7^ A549 cells for each sample were processed using Magna RIP™ RNA-Binding Protein Immunoprecipitation Kit (Millipore Sigma) following the manufacturer’s instructions. AGO2 RIP-grade antibody (Proteintech) and normal mouse IgG (Millipore Sigma) were employed for immunoprecipitation. RNA was purified using UltraPure™ Phenol:Chloroform:Isoamyl Alcohol (25:24:1, v/v) (Thermo Fisher), quantified with Nanodrop 2000c instrument (Thermo Fisher Scientific), and then analyzed by qRT-PCR using specific Taqman probes as explained in the section “RNA isolation, Reverse transcription, and Real-Time RT-PCR.”

A small volume of immunoprecipitated samples (10% of the total) were diluted in Laemmli SDS sample buffer (Thermo Fisher) and loaded in 4- 12% Mini-PROTEAN Tris-Tricine Precast Gels or Criterion Tris-Tricine Precast Gels (Bio-Rad). Anti- AGO-2 (Novus) and Vinculin (Abcam) antibodies were used for western blotting analysis.

The normal mouse IgG sample was conceived as the negative control, while the INPUT sample (10% of the total lysate before immunoprecipitation) was considered the positive one.

### Statistical tests and analyses of publicly available data

Statistical tests used throughout this work are described in the Supplementary Information and indicated in the figure legends where appliable.

The TCGA (v20) and TARGET (v20) miRNA-Seq samples (BAM file format), along with patients’ clinical data, gene raw reads count/FPKM expression data, were retrieved via the Genomic Data Commons Data Portal. See Supplementary Information for more details.

### Data and code availability

The data and source code produced in this work will be stored on Code Ocean and Zenodo repositories. A full description of all methods and reagents can be found in Supplementary Information.

## RESULTS

### MiRNA Isoforms Profiling

We used miRNA-Seq sample data from two large cancer datasets, TCGA and TARGET, to investigate isomiRs at a large scale. The cohorts’ essential characteristics are summarized in Table 1. An in-house workflow (Figure 1A, see Supplementary Information), based on miRge2.0 pipeline (30), was employed to profile canonical miRNA/isomiR molecules, including edited miRNAs (Table S2). On average, we identified 2,511 ±291 (*SD*) expressed molecules (see Supplementary Information) per cohort, detecting nine novels expressed isomiRs. Overall, the number of expressed isomiRs was about *8*-fold the number of expressed canonical miRNAs (miRBase v22). Notably, TCGA-TGCT (*testicular germ cell tumors*) and TCGA-GBM (*glioblastoma multiforme*) cohorts displayed the highest number of expressed isomiRs (Figure 1C).

**Table 1.** TCGA/TARGET Cohorts Basic Characteristics. The table reports cohorts’ essential characteristics, including the number of cases, age at diagnoses, gender, and race.

| **Cohort** | **Cancer type** | **No. of Cases** | **Age at diagnoses**<br>**(Mean ± SD /NA)** | **Gender**<br>**(M/F/NA)** | **Race**<br>**(White/AA/Other/NA)** |
| --- | --- | --- | --- | --- | --- |
| TARGET-ALL-P2 | Acute Lymphoblastic Leukemia - Phase II | 191 | 6.8 ± 5.3 /0 | 101/90/0 | 133/20/7/31 |
| TARGET-ALL-P3 | Acute Lymphoblastic Leukemia - Phase III | 38 | 8.4 ± 5.3 /0 | 21/17/0 | 1/1/0/36 |
| TARGET-AML | Acute Myeloid Leukemia | 701 | 9.1 ± 6.1 /22 | 344/335/22 | 502/77/33/89 |
| TARGET-RT | Rhabdoid tumors | 66 | 1.2 ± 2.2 /0 | 35/31/0 | 49/8/0/9 |
| TARGET-WT | Wilms tumor | 127 | 4.1 ± 2.8 /0 | 53/74/0 | 95/18/0/14 |
| TCGA-ACC | Adrenocortical carcinoma | 80 | 46.4 ± 15.9 /0 | 31/49/0 | 67/1/1/11 |
| TCGA-BLCA | Bladder Urothelial Carcinoma | 409 | 68.0 ± 10.6 /1 | 302/107/0 | 324/23/44/18 |
| TCGA-BRCA | Breast invasive carcinoma | 1079 | 58.6 ± 13.2 /17 | 12/1066/1 | 746/182/62/89 |
| TCGA-CESC | Cervical squamous cell carcinoma and endocervical adenocarcinoma | 307 | 48.2 ± 13.8 /2 | 0/307/0 | 211/30/30/36 |
| TCGA-CHOL | Cholangiocarcinoma | 36 | 63.0 ± 12.8 /0 | 16/20/0 | 31/2/3/0 |
| TCGA-COAD | Colon adenocarcinoma | 444 | 66.8 ± 13.1 /4 | 231/211/2 | 213/59/12/160 |
| TCGA-DLBC | Lymphoid Neoplasm Diffuse Large B-cell Lymphoma | 47 | 56.3 ± 14.1 /0 | 22/25/0 | 28/1/18/0 |
| TCGA-ESCA | Esophageal carcinoma | 184 | 62.5 ± 11.9 /0 | 157/27/0 | 114/5/45/20 |
| TCGA-GBM | Glioblastoma multiforme | 5 | 0 ± 0 /5 | 0/0/5 | 0/0/0/5 |
| TCGA-HNSC | Head and Neck squamous cell carcinoma | 524 | 60.9 ± 11.9 /1 | 383/141/0 | 449/47/13/15 |
| TCGA-KICH | Kidney Chromophobe | 66 | 51.5 ± 14.3 /0 | 39/27/0 | 58/4/2/2 |
| TCGA-KIRC | Kidney renal clear cell carcinoma | 516 | 60.5 ± 12.1 /0 | 335/181/0 | 445/56/8/7 |
| TCGA-KIRP | Kidney renal papillary cell carcinoma | 291 | 61.5 ± 12.1 /5 | 214/77/0 | 207/61/8/15 |
| TCGA-LAML | Acute Myeloid Leukemia | 188 | 54.9 ± 16.2 /0 | 101/87/0 | 171/13/2/2 |
| TCGA-LGG | Brain Lower Grade Glioma | 512 | 43.0 ± 13.4 /2 | 281/230/1 | 471/21/9/11 |
| TCGA-LIHC | Liver hepatocellular carcinoma | 373 | 59.3 ± 13.4 /4 | 254/119/0 | 183/17/163/10 |
| TCGA-LUAD | Lung adenocarcinoma | 513 | 65.3 ± 9.9 /30 | 239/274/0 | 387/52/8/66 |
| TCGA-LUSC | Lung squamous cell carcinoma | 478 | 67.4 ± 8.6 /9 | 354/124/0 | 333/30/9/106 |
| TCGA-MESO | Mesothelioma | 87 | 63.0 ± 9.8 /0 | 71/16/0 | 85/1/1/0 |
| TCGA-OV | Ovarian serous cystadenocarcinoma | 489 | 59.9 ± 11.5 /11 | 0/486/3 | 422/32/18/17 |
| TCGA-PAAD | Pancreatic adenocarcinoma | 178 | 64.6 ± 10.9 /0 | 98/80/0 | 157/6/11/4 |
| TCGA-PCPG | Pheochromocytoma and Paraganglioma | 179 | 47.3 ± 15.1 /0 | 78/101/0 | 148/20/7/4 |
| TCGA-PRAD | Prostate adenocarcinoma | 494 | 61.0 ± 6.8 /11 | 494/0/0 | 146/7/2/339 |
| TCGA-READ | Rectum adenocarcinoma | 161 | 64.2 ± 11.8 /1 | 86/74/1 | 81/6/1/73 |
| TCGA-SARC | Sarcoma | 259 | 60.8 ± 14.7 /1 | 119/140/0 | 227/18/6/8 |
| TCGA-SKCM | Skin Cutaneous Melanoma | 448 | 58.1 ± 15.6 /8 | 276/172/0 | 425/1/12/10 |
| TCGA-STAD | Stomach adenocarcinoma | 436 | 65.7 ± 10.7 /9 | 281/155/0 | 273/13/88/62 |
| TCGA-TGCT | Testicular Germ Cell Tumors | 150 | 32.0 ± 9.3 /16 | 134/0/16 | 119/6/4/21 |
| TCGA-THCA | Thymoma | 506 | 47.3 ± 15.8 /0 | 136/370/0 | 334/27/53/92 |
| TCGA-THYM | Thyroid carcinoma | 124 | 58.2 ± 13.0 /1 | 64/60/0 | 103/6/13/2 |
| TCGA-UCEC | Uterine Carcinosarcoma | 550 | 63.9 ± 11.2 /15 | 0/539/11 | 367/107/33/43 |
| TCGA-UCS | Uterine Corpus Endometrial Carcinoma | 57 | 69.7 ± 9.2 /0 | 0/57/0 | 44/9/3/1 |
| TCGA-UVM | Uveal Melanoma | 80 | 61.6 ± 13.9 /0 | 45/35/0 | 55/0/0/25 |

**Figure 1.**
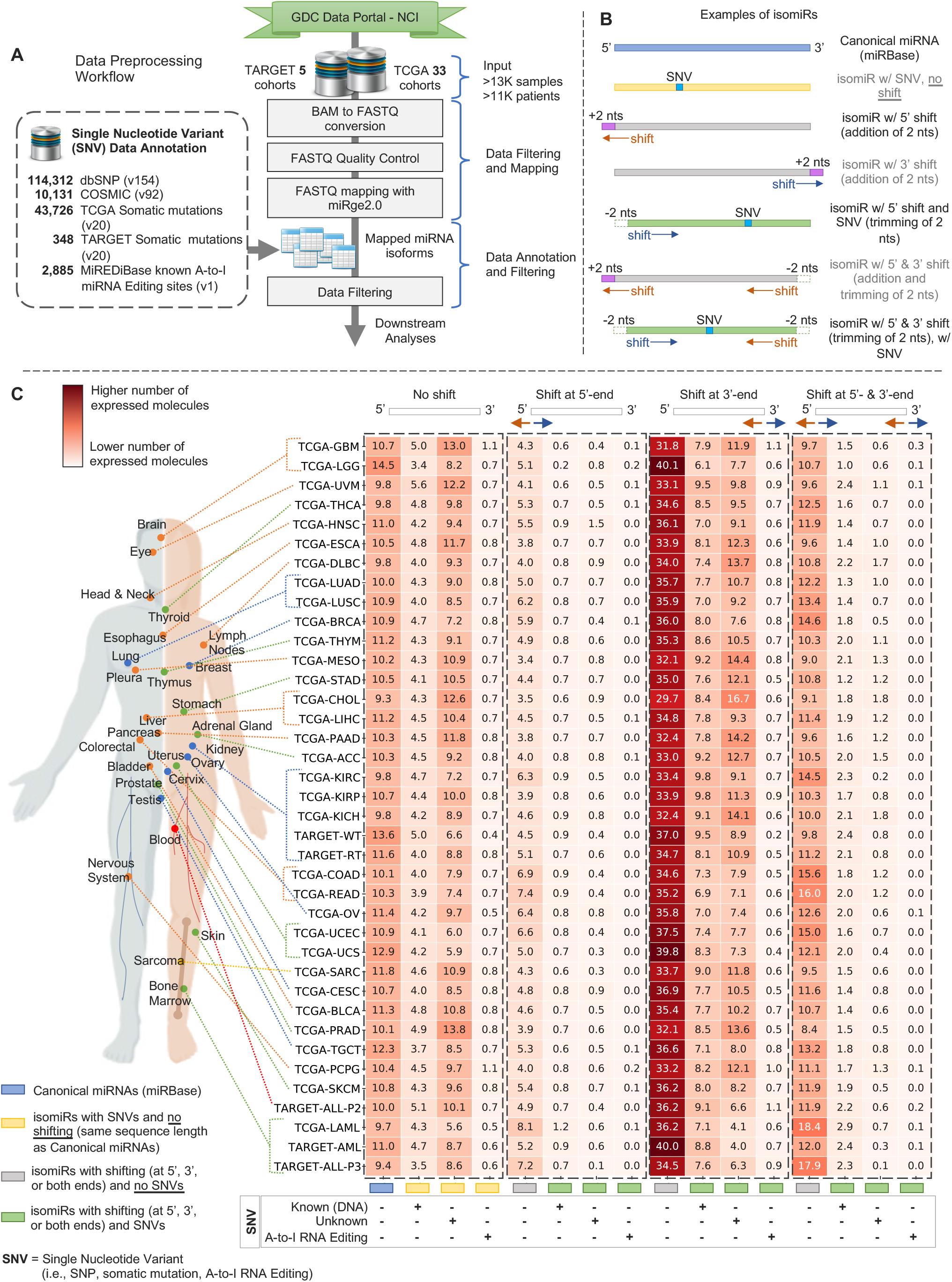
Data Preprocessing Workflow, isomiRs Classification, and Modification Types Distribution. (A-C) In-house data preprocessing workflow (A), examples of annotated isomiRs (B), and distribution of expressed molecules across cohorts and modification types (C). See Table S2 for the complete list of A-to-I RNA editing sites employed by the workflow. See Table S3 for more detailed information regarding the distribution of modification types of expressed molecules across 5p and 3p arms, the shifting amount at 5’- and 3’-ends, along with the number of molecules affected by SNPs, somatic mutations, and A-to-I RNA editing sites. See Supplementary Information for more details.

Of all identified RNA modifications, the 3’-shifted isomiR group was the most abundant (Figure 1C), showing an expression comparable to the canonical one (Figure S1). 3’-shifted isomiRs are highly present in TCGA-TGCT, TCGA-THYM (*thymoma*), TCGA-UCS (*uterine carcinosarcoma*), TCGA-GBM, TCGA-LAML (*acute myeloid leukemia*), TARGET-RT (*rhabdoid tumors*), TCGA-SKCM (*skin cutaneous melanoma*), TCGA-PCPG (*pheochromocytoma and paraganglioma*), and TCGA-ACC (*adrenocortical carcinoma*). The TCGA-LAML showed the highest number of expressed 5’-shifted isomiRs, along with TARGET-ALL-P3 (*acute lymphoblastic leukemia phase 3*), TCGA-READ (*rectum adenocarcinoma*), TCGA-TGCT, TCGA-COAD (*colon adenocarcinoma*), TCGA-UCEC (*uterine corpus endometrial carcinoma*), TCGA-LUSC (*lung squamous cell carcinoma*), TCGA-OV (*ovarian serous cystadenocarcinoma*), and TCGA-THYM. The TCGA-GBM, TCGA-UVM, TCGA-THYM, TCGA-ACC, TARGET-ALL-P2 (*acute lymphoblastic leukemia phase 2*), TCGA-PCPG, TCGA-LAML, TCGA-TGCT, and TARGET-RT cohorts exhibited the highest number of expressed isomiRs with known SNVs (i.e., SNPs and somatic mutations). Besides, the TCGA-GBM, TCGA-PCPG, TARGET-ALL-P2, TCGA-TGCT, TCGA-UVM (*uveal melanoma*), TARGET-ALL-P3 (*acute lymphoblastic leukemia phase 3*), TCGA-ACC, TCGA-SKCM, and TCGA-THCA (*thyroid carcinoma*) were the most enriched cohorts with A-to-I miRNA editing sites (Table S3). Lastly, the TCGA-CHOL (*cholangiocarcinoma*), TCGA-GBM, TCGA-PAAD (*pancreatic adenocarcinoma*), TCGA-MESO (*mesothelioma*), TCGA-ACC, TCGA-PCPG, TCGA-DLBC (*lymphoid neoplasm diffuse large b-cell lymphoma*), TCGA-THYM, and TCGA-ESCA (*esophageal carcinoma*) cohorts displayed the uppermost number of unknown SNVs.

To investigate the potential differences due to Drosha/Dicer cleavage on generating isomiRs across each miRNA arm, we assessed the abundance of modification types (Table S3 - “*Expressed molecules across arms*”). Overall, the 3’-shifted isomiR (no SNVs) represented the predominant modification in both arms. We later looked at the two arms’ 5’ and 3’-end. On average, the 5p arm 5’-end showed the highest stability (Table S3 - “*5p arm - Shifting*”), with ~83% expressed isomiRs characterized by no shifting at all (5’-end untouched, no nucleotide added or trimmed) and ~10% with one nucleotide trimmed (one nucleotide removed at 5’-end). Similarly, the 3p arm 5’-end displayed comparable stability (Table S3 - “*3p arm - Shifting*”), although a bit lower (~75%), promoting the addition (~9%) and trimming (~12%) of one nucleotide at 5’-end, respectively. By stark contrast, the 3’-end revealed higher mobility. The percentage of expressed isomiRs with no 3’-end shifting plunged to ~30% (5p arm) and ~33% (3p arm). In much the same way (Table S3 - “*Arms shifting comparison*”), the two arms consistently showed trimming of 5 (~4%), 4 (~4%), 3 (~6%), 2 (~10%), and 1 (~21%) nucleotides, along with the addition of 1 (~14%) to 2 (~5%) nucleotides. We examined the SNVs distribution along a hypothetical miRNA isoform sequence of ~26 nts long to include the farthest known SNV observed in our expressed molecules. Interestingly, the A-to-G, C-to-T, G-to-A, and T-to-C represented the most predominant SNP forms in both 5p and 3p arms (Table S3 - “*5p and 3p arms – SNPs*”). They were scattered along the sequence length, with the last three mainly located close to the 3’-end. Switching to the 5p arm, the G-to-T, T-to-A, and T-to-G forms were primarily located near the 3’-end, nearby the 21^st^ nucleotide. Aside, the remaining fewer present forms were somehow scattered along the sequence. Notably, the 5p arm was more susceptible to somatic mutations, even though no particular somatic mutation form emerged above the others (Table S3 - “*5p and 3p arms - Somatic mut*.”). A-to-I RNA editing sites were mainly located within the 1-10 nts region (seed region included) in both 5p and 3p arms, with few additional sites involving the 15-24 nts region (Table S3 - “*5p and 3p arms - A-to-I RNA Edit*.”). Altogether, the distribution of modification types was somehow balanced between 5p and 3p arms, with the 5p arm leading by up to 1% more expressed molecules over the 3p arm.

Finally, to gain insights into the mechanisms that may lead to the isomiRs accumulation in cancer, we explored the differences in the abundance of expressed isomiRs across cohorts’ cancer samples. For each cohort/cancer tissue, we compared samples characterized by a low (first quartile) against a high (third quartile) number of expressed isomiRs. Here, we retained significantly dysregulated genes (see Supplementary Information). We then performed a pathways enrichment analysis via Ingenuity^®^ Pathway Analysis (IPA) software (Table S4; see Supplementary Information). Finally, in Figure S2, we reported the most significant pathways enriched in at least five cohorts/cancer tissues. The difference in the abundance of expressed miRNA isoforms between the two groups, low and high, characterized the activation/deactivation of several critical pathways involved in proliferation, metastasization, tumor immune escape, invasion, and angiogenesis, such as the ILK, HIF1α, and Rac signaling pathways, PD-1/PD-L1 cancer immunotherapy pathway, and regulation of the epithelial-mesenchymal transition (EMT) by growth factors pathway.

### MiRNA Isoforms-based Clustering Reveals Unique Clinical-Pathological Stratification

Purely for clustering purposes, we benchmarked three sets of expressed molecules grouped according to their modification type (Figure S3A; see Supplementary Information). We investigated the benefits and drawbacks of using specific sets of molecules by assessing their ability to cluster samples across different cohorts/cancer tissues. In the first set, labeled “*CAN*,” we considered only canonical miRNAs (miRBase v22). In the second one, marked “*ISO_wo_SNV*,” we used both canonical miRNAs and shifted isomiRs without SNVs. In the last set, labeled “*ISO*,” we employed all expressed canonical miRNAs and isomiRs, including the shifted ones.

We applied an in-house designed workflow to each of the three sets (Figure S3B; see Supplementary Information). By benchmarking the three sets, we observed a general trend in which the *ISO* set reached a higher clustering ability (Figure 2A). The *ISO*-based clustering was able to better separate cohorts’ cancer samples than the other two sets (Figure 2B). We then examined clustering results from a clinical-pathological perspective, focusing solely on available and most significant clinical-pathological features (*Chi-Square p-value* < 0.01) (Table S5). Overall, the *CAN-*, *ISO_wo_SNV-*, and *ISO*-based findings somehow supported one another in clustering results (Figure S4A-C). For instance, in the TCGA-LIHC and TCGA-TGCT cohorts, both *CAN* and *ISO_wo_SNV* sets grouped patients according to their clinical stages (Figure S4A-B). At the same time, both *CAN* and *ISO* sets significantly clustered patients in the TCGA-STAD cohort, though reflecting different clinical-pathological features (Figure S4A, S4C).

**Figure 2.**
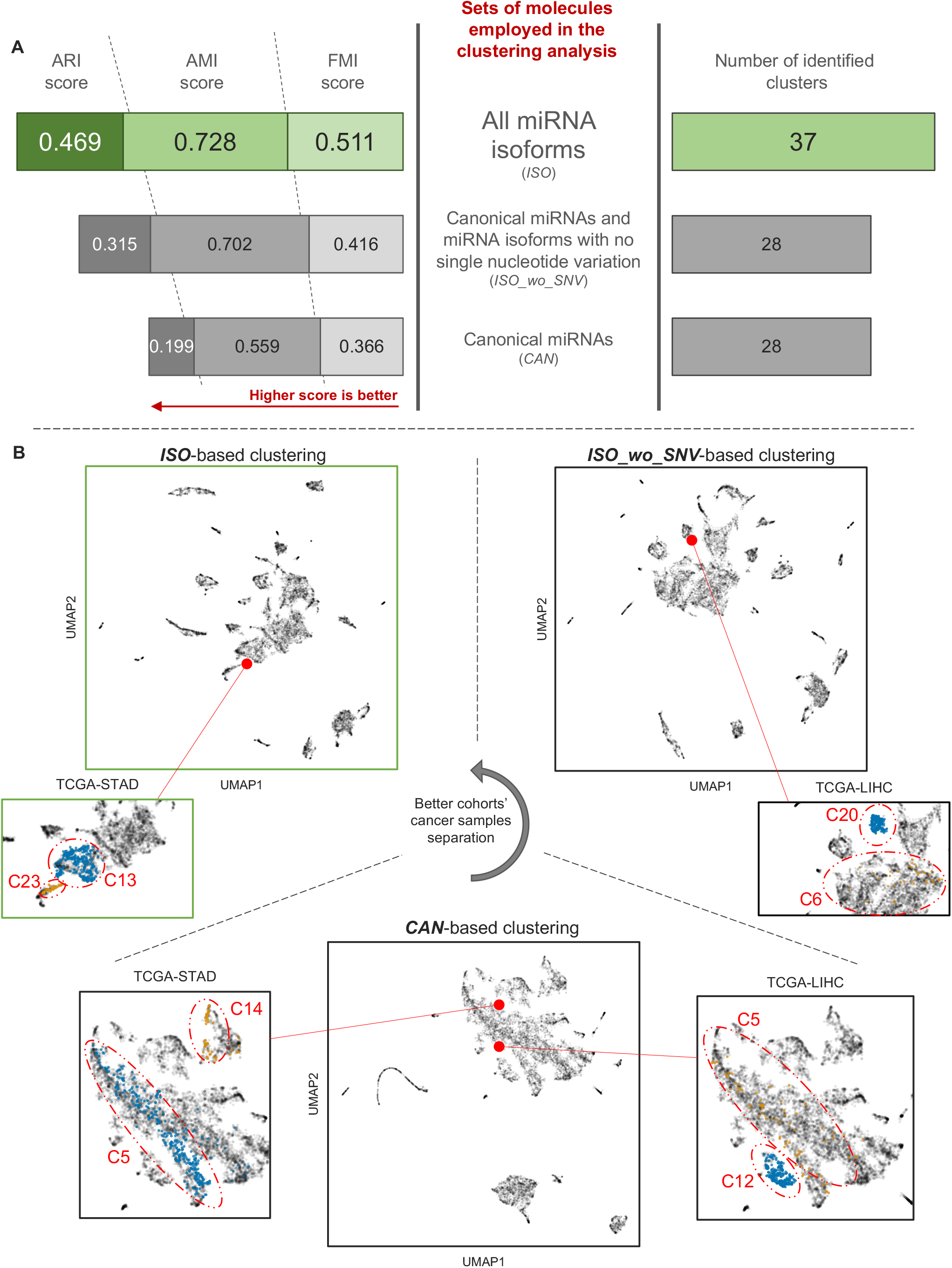
MiRNA Isoform-based Clustering Better Delineates Clinical-Pathological Stratification. (A-B) Clustering benchmarks results related to three different sets (*CAN*, *ISO_wo_SNV*, and *ISO*) of molecules (A) and a comparison between *the three* sets to highlight their ability to separate cohorts’ cancer samples (B). Panel A reports quality scores (*Adjusted Rand Index* - ARI, *Adjusted Mutual Information* - AMI, and *Fowlkes-Mallows Index* - FMI) and the number of identified clusters for each set. Panel B compares clustering based on *CAN*, *ISO_wo_SNV*, and *ISO* sets, highlighting cancer sample separation. See Figure S2A-B for more detailed information on how we defined the three sets (A) and the designed workflow (B) used for data visualization and benchmarking. See Figure S3A-C for a complete comparison between *CAN*-, *ISO_wo_SNV-*, and *ISO*-based clustering. See Table S5 for more detailed information on the most prominent and significant clinical-pathological features taken into account for clustering-based clinical-pathological analysis. See Supplementary Information for more details.

By including miRNA isoforms data (*ISO_wo_SNV* and *ISO* sets), we obtained a more refined classifier, which highlighted, in some cohorts, additional subclusters with clinical-pathological relevance. Furthermore, unlike the *CAN* set, both *ISO_wo_SNV* and *ISO* set aggregated the TCGA-ESCA and TARGET-AML cancer samples in the same way. In TCGA-ESCA, patients were split according to their histological type, squamous (*C6* in *ISO_wo_SNV*, *C12* in *ISO*) and adeno (*C11* in *ISO_wo_SNV*, *C13* in *ISO*). In TARGET-AML, cancer samples were grouped in two clusters, *C0* and *C3*, consistently with their cytogenetic complexity, a well-known prognostic marker (Figure S4B-C; Table S5). Notably, besides having a lower cytogenetic complexity, cluster *C0* included cancer samples that harbored chromosomal translocations commonly associated with good prognosis, t(9;11)(p22;q23) and inv(16) (31,32). In stark contrast, cluster *C3* was enriched with FLT3-ITD positive samples associated with poor survival (33).

Finally, yet importantly, the three sets were able to cluster cohorts’ cancer samples exclusively. The *CAN* set uniquely partitioned cancer samples in TCGA-HNSC, distinguishing among patients graded as well (G1) (cluster *C17*) and poorly (G3) (*cluster C5*) differentiated (Figure S3A). According to the clinical stage, the ISO_wo_SNV set separated patients in the TCGA-KIRP cohort (Figure S4B). Furthermore, the *ISO* set clustered the TCGA-COAD and TCGA-READ cohorts’ patients into two groups, in which cluster C23 was characterized by lymphatic invasion and the presence of a history of polyps. In TCGA-LUSC, patients were grouped according to their clinical stages (Figure S4C; Table S5).

### Differentially Abundant miRNA Isoforms Across Cancer Tissues

To determine whether miRNA isoforms were dysregulated across cohorts/cancer tissues, we performed a differential expression (DE) analysis comparing *primary solid*, *recurrent solid*, *metastatic*, and *normal tissues* (see Supplementary Information). The most significantly dysregulated molecules were retained according to an *adjusted p-value* <0.05 and |*linear fold change*| >1.5 (Table S6). The resulting molecules were characterized by a similar trend outlined in the “*MiRNA Isoforms Profiling*” paragraph (Figure 1B). They were affected mainly by 3’-end shifting in almost all cohorts/comparisons, followed by miRNA isoforms with both 5’- and 3’-end shifting. Distributions of dysregulated molecules per cohort/comparison over isomiRs modification types are reported in Figure 3A, including four novel dysregulated miRNA isoforms. We then examined the modification types distribution across 5p and 3p arms (Table S7 - “*Mod. types distribution - Arms*”). In most cohorts/comparisons, the 5p arm showed ~1.12% ±0.46 (*SD*) additional molecules than the other arm. Once again, the 3’-end shifting modification (no SNVs) exhibited the highest number of dysregulated molecules in both arms. Considering the subtle difference between the 5p and 3p arms, we aggregated the two arms’ contribution, reporting the most noticeable results in Figure 3B. The 5’-end was confirmed to be the most stable of the two ends, with ~70-84% dysregulated molecules characterized by no 5’-end shifting, in sharp contrast to the 3’-end, whose stability sank to ~27-37% (Figure 3B; Table S7 - “*5’- and 3’-end shifting*”). The 5’-end was characterized by one nucleotide added and trimmed, while the 3’-end displayed a broader shifting, trimming 1 to 5 nts and adding 1 to 2 nts (Figure 3B; Table S7 - “*5’- and 3’-end shifting*”).

**Figure 3.**
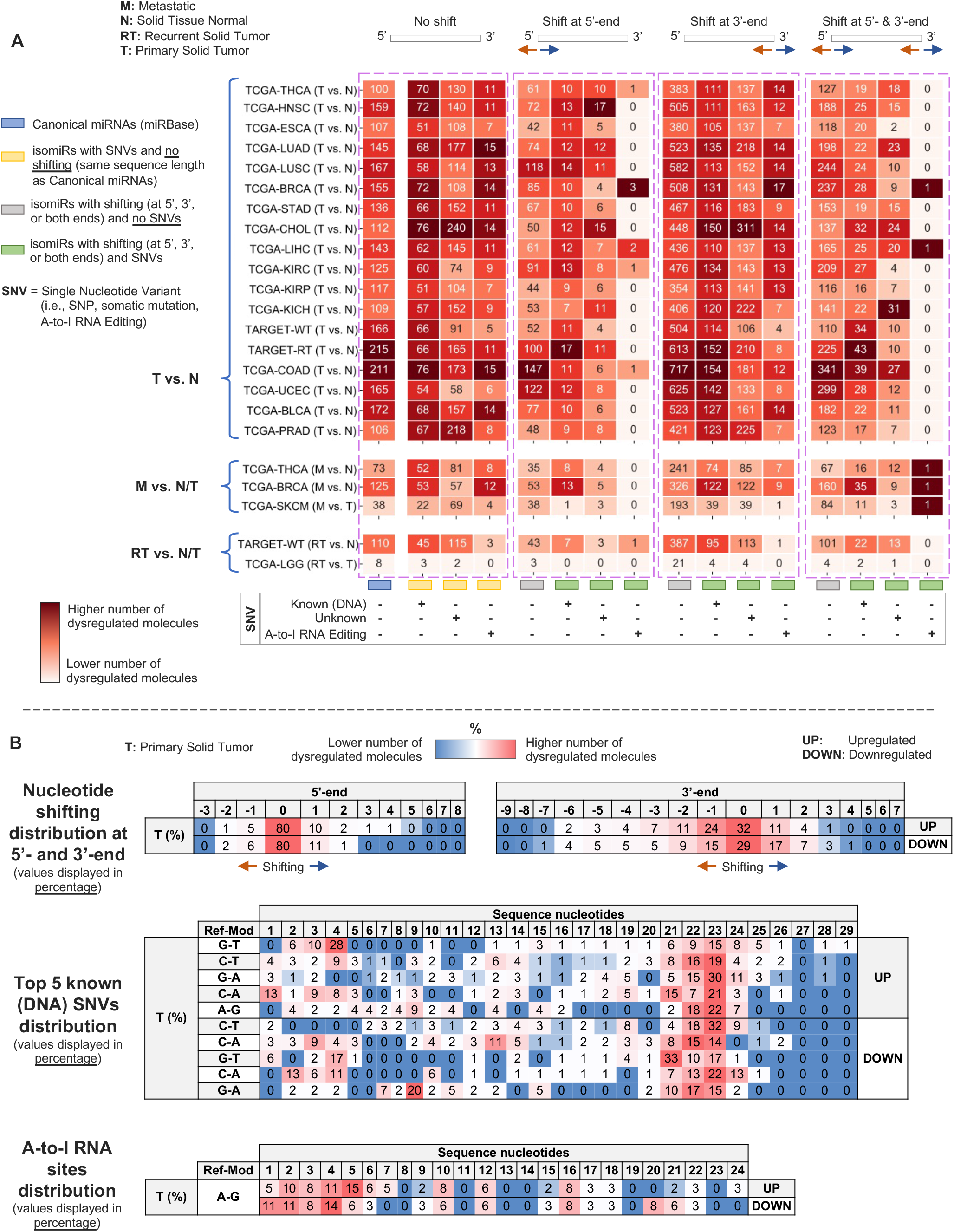
MiRNA Isoforms Dysregulated Across Cohorts and Tissues. (A-B) Distribution of dysregulated molecules per cohort/comparison and modification type (A), and most prominent modification types (5’- and 3’-end shifting, SNPs/somatic mutations, and A-to-I RNA editing sites) (B). See Table S6 for the complete information on dysregulated molecules across cohorts/comparisons, and Table S7 for detailed information on all modification types affecting dysregulated molecules (5’- and 3’-end shifting, SNPs/somatic mutations, and A-to-I RNA editing sites). See Supplementary Information for more details.

Moving the attention to known DNA SNVs, we identified the G-to-T, C-to-T, and G-to-A forms to be the most frequent variants affecting upregulated molecules, whereas downregulated molecules mainly faced G-to-A, C-to-T, and C-to-A modifications (Figure 3B; Table S7 - “*SNPs-Somatic mut. distribution*”). Furthermore, these known DNA SNVs were spread along the sequence length in both upregulated and downregulated molecules. Finally, A-to-I RNA editing sites were still primarily located within the 1-10 nts region (seed region included), with few exceptions in the 15-24 nts region (Figure 3B; Table S7 - “*A-to-I RNA Edit. distribution*”).

### Dysregulated Canonical miRNAs and IsomiRs with Opposite Expression Trend in Cancer Reveal Different Behavior

Taking a closer look at the dysregulated canonical miRNAs across cohorts, we observed 104 out of 573 canonical miRNAs being characterized by an opposite expression trend compared to their miRNA isoforms (Table S8). In this regard, we searched for a candidate among the 104 canonical miRNAs to assess the potential gene targeting shifting between miRNA isoforms and their canonical counterparts. Interestingly, the canonical miR-101-3p appeared to be downregulated in 6 cohorts (TCGA-LUAD, TCGA-LIHC, TCGA-HNSC, TARGET-RT, TARGET-WT, and TCGA-CHOL), with the sole TCGA-LUAD cohort reporting miRNA isoforms lacking SNVs, also confirmed by a previous study (34). Therefore, we elected to investigate the canonical miR-101-3p and one of its isomiRs in lung adenocarcinoma (TCGA-LUAD), comparing *primary solid* against *normal tissue*. Our interest in investigating such a canonical miRNA was corroborated by a recent work in which authors assessed one isomiR of canonical miR-101-3p in the human brain (35). Furthermore, the authors demonstrated the isomiR ability to negatively modulate the expression of five validated miR-101-3p targets, leading them to consider the isomiR as a miR-101-3p functional variant.

In our work, we studied one isomiR, miR-101-3p (−1|-2), characterized by one nucleotide added at 5’-end, termed “-1,” and two nucleotides trimmed at 3’-end, termed “−2” (see Supplementary Information). As shown in Figures 4A and 4F, the two molecules were characterized by an opposite expression trend, with the canonical miRNA resulting downregulated in cancer samples. Looking for putative and exclusive targets, we retained dysregulated genes (Table S9) based on their significance (see Supplementary Information) and opposite expression trends (i.e., miRNA *up* and genes *down*, or vice versa) (Figure 4B, 4G). Finally, the set of dysregulated targets was intersected with the list of predicted gene targets generated by *isoTar* (36) (Table S9), requiring a minimum consensus of two prediction tools. Out of the reduced set of genes, we elected *Prostaglandin-Endoperoxide Synthase 2* (*PTGS2* or *COX-2*), a gene studied in cancers (37,38), which is a validated target for miR-101-3p and overexpressed in lung cancer (39), and *Desmocollin-2* (*DSC2*), a putative target for the miR-101-3p (−1|−2) with a low expression in lung cancer (40). PTGS2 promotes tumor growth, angiogenesis, and tissue invasion. It also induces resistance to therapeutic agents, compromising tumor immunity and apoptosis (41). In line with the literature, we demonstrated the direct binding (Figure S5A) through luciferase assay (Figure 4C) between this oncogene and the downregulated canonical miR-101-3p. After miR-101-3p ectopic overexpression in HEK-293 cells, we observed ~40% reduction of luciferase activity compared to the scramble negative control (SCR) (Figure 4C). On the other hand, overexpression of miR-101-3p (−1|−2) resulted in a minor (20%) reduction of luciferase activity (Figure 4C). After transfecting the canonical miR-101-3p (Figure S5B), western blotting experiments highlighted a significant downregulation of endogenous PTGS2 in the A549 and H1299 lung cancer cell lines, while no variation of PTGS2 was observed transfecting miR-101-3p (−1|−2) (Figure 4D, 4E). Our findings demonstrated that of the two molecules, only the downregulated canonical miR-101-3p exclusively targeted PTGS2, which in turn is upregulated in lung cancer.

**Figure 4.**
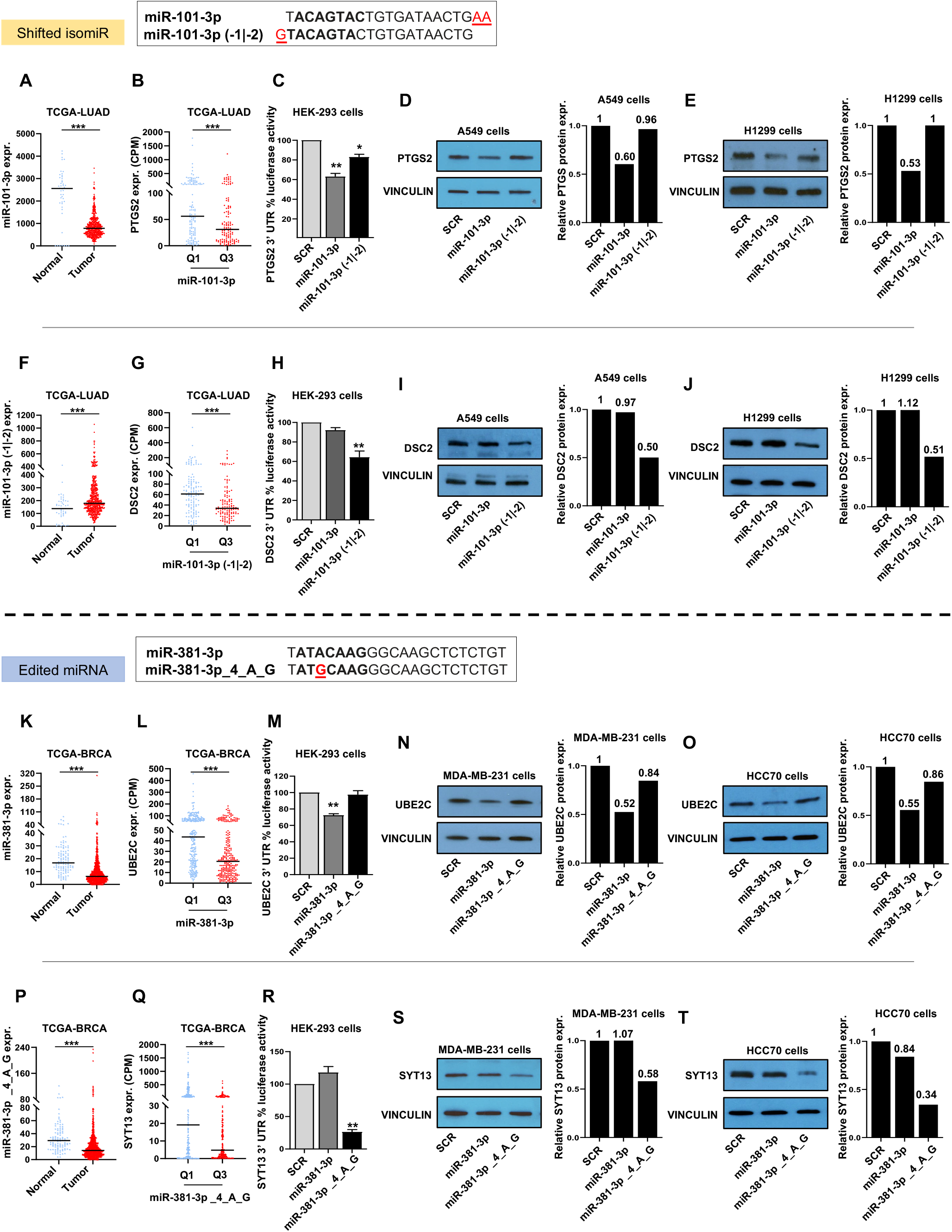
MiRNA Isoforms Experimental Gene Targeting Validation. (A-E) *miR-101-3p* and (F-J) *miR-101-3p (−1|-2)* experimental targeting validation in lung cancer cells. Expression of miR-101-3p (A) and miR-101-3p (−1|-2) (F) in normal and tumor samples in TCGA-LUAD cohort. PTGS2 (B) and DSC2 (G) expression in TCGA-LUAD samples in miR-101-3p/miR-101-3p (−1|-2) first (Q1) and third (Q3) quartile. Luciferase assay for psiCHECK-2-PTGS2 3’ UTR WT (C) and psiCHECK-2-DSC2 3’ UTR WT (H) constructs co-transfected with mirVana™ miRNA mimics for miR-101-3p, miR-101-3p (−1|-2), and negative scramble miRNA control (SCR) in HEK-293 cells performed 24 hours after the transfection. Western blotting depicts the downregulation of PTGS2 (D-E) or DSC2 (I-J) proteins in A549 and H1299 cells, respectively, after miR-101-3p and miR-101-3p (−1|-2) overexpression. Densitometric quantification of western blotting signals (D-E, I-J) was performed using ImageJ (U. S. National Institutes of Health, Bethesda, Maryland, USA, https://imagej.nih.gov/ij/, 1997– 2018). (K-O) *miR-381-3p* and (P-T) *miR-381-3p_4_A_G* experimental targeting validation in Triple-Negative (TN) breast cancer cells. Expression of both miR-381-3p miRNA isoforms in normal and breast cancer samples in TCGA-BRCA cohort (K, P). Luciferase assay for psiCHECK-2-UBE2C 3’ UTR WT (M) and psiCHECK-2-SYT13 3’ UTR WT (R) constructs co-transfected with mirVana™ miRNA mimics for miR-381-3p, miR-381-3p_4_A_G, and negative scramble miRNA control (SCR) in HEK-293 cells performed 24 hours after the transfection. Western blotting represents the downregulation of UBE2C (N-O) and SYT13 (S-T) proteins in MDA-MB-231 and HCC70 cells after miR-381-3p and miR-381-3p_4_A_G upregulation via mirVana miRNA mimic transfection. The histogram reports densitometric quantification of western blotting signals (N-O, S-T), were performed using ImageJ (U. S. National Institutes of Health, Bethesda, Maryland, USA, https://imagej.nih.gov/ij/, 1997–2018). Pictures are representative of at least three experiments. The fold of increase in the graphics is the mean values of 3 replicates. *P-value* <0.05 was considered statistically significant. Annotations for * 0.01 ≤ *p-value* <0.05, ** 0.001 ≤ *p-value* <0.01, and *** *p-value* <0.001 are provided accordingly. Error bars indicate the standard deviation (SD) for the three biological replicates. See Table S9 for more details.

DCS2 is a protein implicated in mediating cell adhesion and epithelial cell proliferation, as well as tumorigenesis (42). The loss of DSC2 increases tumor progression through cell proliferation and metastasis (42). As described above for PTGS2, we demonstrated through luciferase assay (Figure 4H) and western blotting (Figure 4I, 4J) the exclusive targeting of DSC2 by miR-101-3p (−1|−2). Even though the two miRNA molecules rise from the same locus, they show different behavior. In fact, in stark contrast with the shifted isomiR, these results corroborated the tumor suppressor role of the canonical miR-101-3p and the oncogenic role of miR-101-3p (−1|−2) in lung cancer.

### Dysregulated A-to-I Edited miRNA Isoforms in Cancer

In addition to investigating shifted isomiRs, we measured the A-to-I RNA editing abundance across cancer cohorts/comparisons, detecting 169 unique dysregulated A-to-I edited miRNA isoforms (Table S6) that originated from 43 distinct miRNA arms. Looking closely, the edited miR-381-3p (A-to-I RNA editing at position 4) resulted in one of the most diffused dysregulated molecules among the before-mentioned ones. Its downregulation interested 11 out of 22 cohorts/comparisons, a trend confirmed by previous studies (25,26) and observed in several tumors, including breast cancer (43).

This work examined the canonical miR-381-3p and one of its edited forms in the breast cancer cohort (TCGA-BRCA), miR-381-3p_4_A_G. The expression of the two molecules exhibited a significant downregulation in cancer samples (Figures 4K and 4P). In line with the previous section, we applied a similar workflow (see Supplementary Information) to assess potential target variability between the two molecules. After retaining significantly dysregulated genes (Table S9) characterized by an opposite expression trend (miRNA *down*, genes *up*) (Figures 4L and 4Q), we crossed them with the list of gene targets predicted by *isoTar* (Table S9). Out of the reduced set of potential direct targets for miR-381-3p, we elected to study *Ubiquitin Conjugating Enzyme E2 C* (*UBE2C*), a gene that promotes breast cancer proliferation, migration, and invasion, and whose overexpression correlates with poor clinical outcomes (44). Using luciferase reporter vectors containing the 3’ UTR of the gene and the two miRNA molecules (canonical and edited one) in HEK-293 cells, we demonstrated the direct binding (Figure S5A) between UBE2C and miR-381-3p (Figure 4M). Following the miR-381-3p overexpression (Figure S5B), western blotting experiments in TNBC cell lines MDA-MB-231 and HCC70 corroborated our findings, demonstrating a significant downregulation of UBE2C (~50%), as depicted by densitometry (Figure 4N, 4O). At the same time, among miR-381-3p_4_A_G targets, we selected *Synaptotagmin 13* (*SYT13*), an oncogene involved in different cancers (45,46). We validated the direct binding (Figure S5A) between SYT13 and miR-381-3p_4_A_G through luciferase assay in HEK-293 cells, with a reduced luciferase activity of ~80% (Figure 4R). After miR-381-3p_4_A_G overexpression (Figure S5B), western blotting experiments in MDA-MB-231 and HCC70 cell lines confirmed the downregulation of SYT13 as shown by densitometry (Figure 4S, 4T).

Once again, our results pointed out the importance of not limiting studies solely to canonical miRNAs, as outlined by the edited miRNA’s ability to target one oncogene exclusively.

### Prognostic miRNA Isoform Signature

In order to estimate each cohort/cancer tissue’s best performing prognostic signature for Overall Survival (OS) and Relapse Free Survival (RFS), we designed an in-house 2-stages workflow (Figure S6; see Supplementary Information). Cohorts’ clinical characteristics are summarized in Table 2.

**Table 2.** TCGA/TARGET Cohorts Clinical Characteristics. The table shows cohorts’ clinical characteristics for both Overall Survival (OS) and Relapse Free Survival (RFS), including the number of cases, stages, number of events/no events (censored).

| Cohort | # Cases | Stages (I/II/III/IV/NA) | Overall Survival (OS) |  |  | Relapse-Free Survival (RFS) |  |  |
| --- | --- | --- | --- | --- | --- | --- | --- | --- |
|  |  |  | # events | # censored | Median Follow-UP (Months) | # events | # censored | Median Follow-UP (Months) |
| TARGET-ALL-P2 | 191 | 0/0/0/0/191 | 80 | 68 | 56.05 | 0 | 0 | 0.00 |
| TARGET-ALL-P3 | 38 | 0/0/0/0/38 | 12 | 17 | 29.33 | 0 | 0 | 0.00 |
| TARGET-AML | 701 | 0/0/0/0/701 | 265 | 413 | 59.30 | 0 | 0 | 0.00 |
| TARGET-RT | 66 | 2/13/27/0/24 | 32 | 26 | 7.98 | 0 | 0 | 0.00 |
| TARGET-WT | 127 | 17/49/41/14/6 | 52 | 75 | 54.27 | 0 | 0 | 0.00 |
| TCGA-ACC | 80 | 9/37/16/16/2 | 29 | 51 | 39.42 | 36 | 39 | 27.40 |
| TCGA-BLCA | 409 | 2/131/139/135/2 | 179 | 229 | 17.87 | 79 | 232 | 16.50 |
| TCGA-BRCA | 1079 | 182/609/244/20/24 | 149 | 929 | 27.57 | 31 | 363 | 33.27 |
| TCGA-CESC | 307 | 163/70/46/21/7 | 71 | 236 | 21.27 | 26 | 172 | 21.12 |
| TCGA-CHOL | 36 | 19/9/1/7/0 | 18 | 18 | 21.50 | 17 | 13 | 10.72 |
| TCGA-COAD | 444 | 73/168/125/65/13 | 101 | 340 | 22.30 | 30 | 1 | 21.80 |
| TCGA-DLBC | 47 | 7/17/5/12/6 | 9 | 38 | 26.37 | 6 | 21 | 26.37 |
| TCGA-ESCA | 184 | 18/82/62/16/6 | 77 | 107 | 13.35 | 43 | 126 | 12.60 |
| TCGA-GBM | 5 | 0/0/0/0/5 | 0 | 0 | 0.00 | 0 | 0 | 0.00 |
| TCGA-HNSC | 524 | 27/85/92/320/0 | 223 | 300 | 21.50 | 64 | 107 | 23.77 |
| TCGA-KICH | 66 | 21/25/14/6/0 | 9 | 56 | 74.93 | 10 | 54 | 73.67 |
| TCGA-KIRC | 516 | 253/55/123/82/3 | 172 | 344 | 39.38 | 2 | 32 | 13.60 |
| TCGA-KIRP | 291 | 180/25/52/16/18 | 44 | 246 | 25.62 | 18 | 140 | 20.40 |
| TCGA-LAML | 188 | 0/0/0/0/188 | 114 | 63 | 12.17 | 0 | 0 | 0.00 |
| TCGA-LGG | 512 | 0/0/0/0/512 | 124 | 385 | 22.60 | 65 | 186 | 20.13 |
| TCGA-LIHC | 373 | 173/86/85/5/24 | 129 | 243 | 19.83 | 102 | 174 | 13.78 |
| TCGA-LUAD | 513 | 277/121/84/24/7 | 182 | 322 | 21.88 | 31 | 156 | 18.40 |
| TCGA-LUSC | 478 | 230/158/80/6/4 | 199 | 273 | 22.25 | 30 | 139 | 17.00 |
| TCGA-MESO | 87 | 10/16/45/16/0 | 73 | 13 | 17.10 | 48 | 32 | 11.80 |
| TCGA-OV | 489 | 1/27/374/80/7 | 308 | 177 | 34.87 | 60 | 0 | 18.43 |
| TCGA-PAAD | 178 | 21/147/3/4/3 | 93 | 85 | 15.48 | 56 | 106 | 13.08 |
| TCGA-PCPG | 179 | 0/0/0/0/179 | 6 | 173 | 25.17 | 15 | 163 | 23.48 |
| TCGA-PRAD | 494 | 0/0/0/0/494 | 10 | 484 | 30.80 | 60 | 44 | 24.87 |
| TCGA-READ | 161 | 29/48/50/24/10 | 26 | 134 | 20.58 | 10 | 0 | 27.15 |
| TCGA-SARC | 259 | 0/0/0/0/259 | 98 | 161 | 31.57 | 91 | 141 | 22.52 |
| TCGA-SKCM | 448 | 74/128/164/23/59 | 210 | 229 | 37.47 | 221 | 215 | 27.58 |
| TCGA-STAD | 436 | 58/128/180/43/27 | 168 | 263 | 14.07 | 45 | 177 | 13.42 |
| TCGA-TGCT | 150 | 59/14/14/0/63 | 4 | 130 | 42.03 | 30 | 99 | 28.80 |
| TCGA-THCA | 506 | 284/52/113/55/2 | 16 | 490 | 31.50 | 25 | 355 | 33.98 |
| TCGA-THYM | 124 | 0/0/0/0/124 | 9 | 114 | 41.77 | 16 | 107 | 38.13 |
| TCGA-UCEC | 550 | 339/49/123/28/11 | 88 | 450 | 30.32 | 28 | 165 | 34.47 |
| TCGA-UCS | 57 | 22/5/20/10/0 | 35 | 22 | 20.37 | 29 | 25 | 12.97 |
| TCGA-UVM | 80 | 0/39/37/4/0 | 23 | 57 | 26.13 | 17 | 62 | 22.33 |

We benchmarked two distinct sets of expressed molecules: canonical miRNAs and miRNA isoforms (canonical ones included). Out of the 26 cohorts/cancer tissues examined, the workflow identified 4 (OS)/3 (RFS) and 9 (OS)/8 (RFS) significant prognostic signatures (*Log Rank Test-based p-value* <0.01 and *AUC* ≥0.7) using canonical miRNAs and miRNA isoforms, respectively (Figure 5; Table S10). Notably, both sets of canonical miRNAs and miRNA isoforms identified OS signatures for TARGET-ALL-P2, TCGA-UVM, and TCGA-ACC cohorts, with the latter also providing RFS signatures.

**Figure 5.**
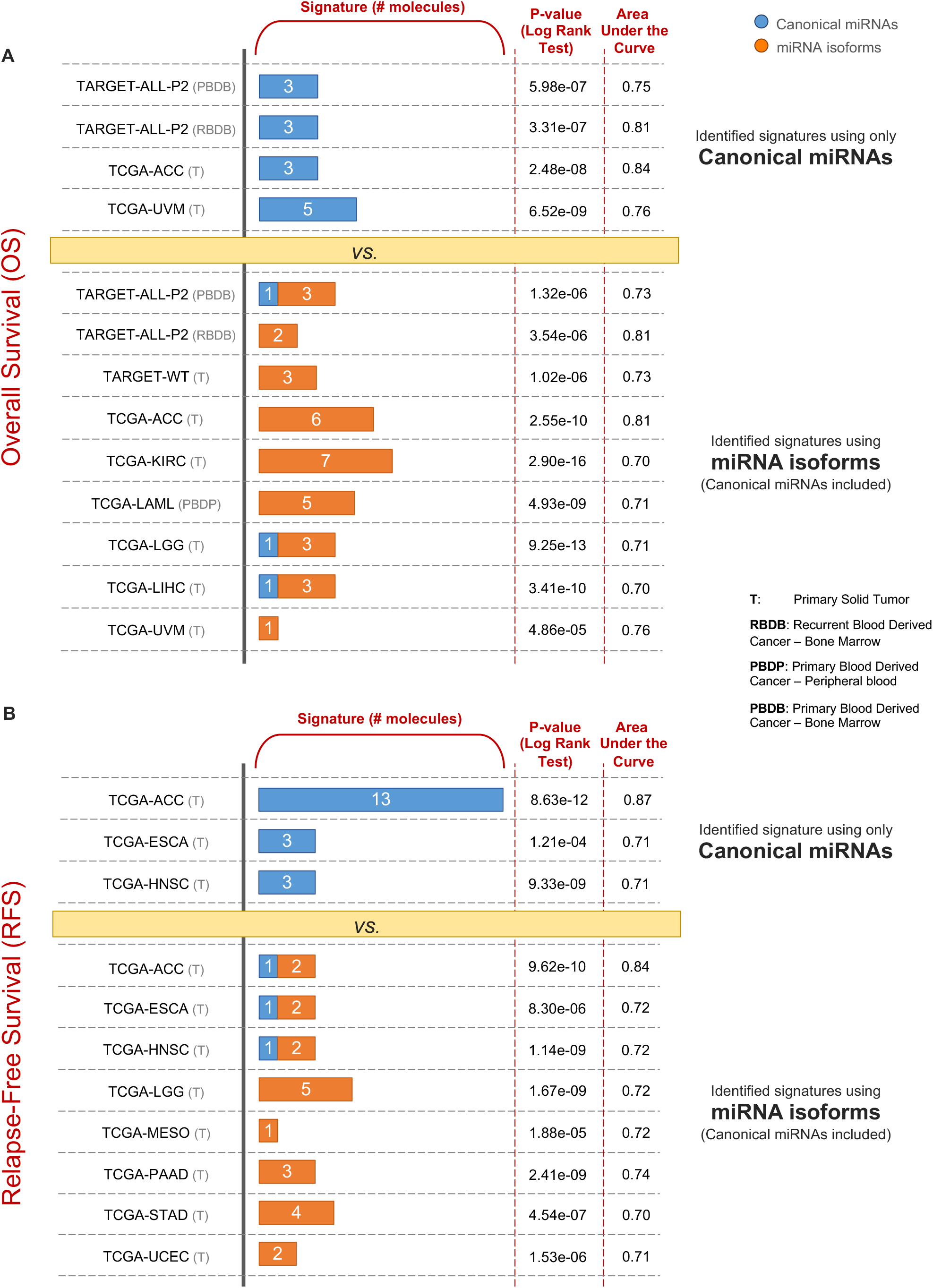
Overall and Relapse Free Survival Risk Score-Based Signature. (A-B) Overview of risk score-based signatures for Overall Survival (A) and Relapse Free Survival (B), supplied with the number of molecules in each signature, the corresponding p-value (Log Rank test), and the area under the curve (AUC) score. See Figure S6 and Table S10 for more detailed information regarding the workflow employed for results generation and the list of molecules for each signature, respectively.

Although both sets produced OS signatures for common cohorts, signatures were characterized by unique molecules. Notably, miRNA isoforms-based OS signatures in TARGET-ALL-P2 (PBDB) and TCGA-KIRC cohorts included a common miRNA isoform of has-miR-26b-3p, characterized by one nt added at 3’-end. By contrast, the sole TCGA-ACC RFS signatures generated using the two sets shared a common canonical miRNA, hsa-let-7c-3p.

Of all identified signatures using the miRNA isoforms set, three signatures for each workflow, OS and RFS, included at least one canonical miRNAs. The OS signatures included the canonical miRNAs hsa-miR-1275, hsa-miR-346, and hsa-miR-941, respectively, in TARGET-ALL-P2 (PBDB), TCGA-LGG, and TCGA-LIHC. The RFS signatures included the canonical miRNAs hsa-let-7c-3p, hsa-miR-503-5p, and hsa-miR-455-5p, respectively, in TCGA-ACC, TCGA-ESCA, and TCGA-HNSC. The complete list of identified signatures, their miRNA isoforms, and additional details are reported in Table S10.

## DISCUSSION

Several scientific contributions have magnified our understanding of the so-called miRNA Epitranscriptome, inflating the interest in RNA modifications such as SNPs (47), A-to-I RNA editing (8,9,25,26,48–50), as well as shifted isomiRs (27–29,51,52), a product of the imprecise miRNA sequence cleavage (10,21,53,54). Although the studies mentioned above have examined miRNA modifications individually, the concurrent occurrence of two miRNA modification categories, A-to-I miRNA editing and shifted isomiRs, has yet to be adequately explored.

In this work, we simultaneously estimated the abundance and implications of a broader set of RNA modifications, processing data at a large scale from the most prominent and reliable public resources, TCGA and TARGET, in a pan-cancer setting. Here, we profiled >13K adult and pediatric cancer samples spread across 38 distinct cohorts. At first glance, several miRNA isoforms displayed a higher expression than their canonical miRNAs, which are the reference molecules in biological databases such as miRBase (v22). In particular, the abundance of expressed miRNA isoforms exceeded by *8*-fold the number of expressed canonical miRNAs. A closer look at the distribution of modification types among expressed miRNA isoforms outlined a predominance of 3’-end shifts, which equally impacted both 5p and 3p arms due to potentially no differences in Drosha and Dicer cleavage. Affected by the addition/trimming of two or more nucleotides, the 3’-end showed higher mobility than the more conservative 5’-end (11). Interestingly, the lower presence of more extended additions (addition of three or more nts at 3’-end) could be explained via degradation processes carried out by some enzymes that remove the exceeding part spurting out the RISC complex (55). As is well known, both 5’- and 3’-end fulfill different functionalities. The first carries out the miRNA-mRNA partial base-pairing through the MSR, though the 3’-end is proven to be critical for the miRNA-mRNA interaction stabilization (18,56,57), especially in the presence of mismatches or bubbles (58). In light of this, the main reason for the high number of expressed 3’-end shifted molecules could be explained as the cell’s attempt to modulate miRNAs activity, perhaps trying to overcome the weakness of specific miRNA-mRNA bindings under particular conditions (58). Likewise, the 5’-end shifting could be the effort of replacing missing canonical miRNAs or the necessity for a targetome shifting (59). Interestingly, more than 40% of dysregulated edited miRNA isoforms reported at least one A-to-I editing site within the MSR. These observations may indicate the tendency of the A-to-I RNA editing phenomenon to give rise to dysregulated miRNA isoforms in cancer, which could exhibit a different targetome and biological function with respect to their canonical counterpart (9). Besides, the distribution of the most representative known DNA variant forms (top-five) observed across dysregulated miRNA isoforms mostly fall near the 3’-end (between the 21^st^ and 23^rd^ nucleotides), potentially impinging either the miRNA lifespan or the targeting stability (55,60). Lastly, from a functional standpoint, we explored the underlying differences in the abundance of expressed miRNA isoforms in each cohort/cancer tissue. We compared cancer samples characterized by a lower (first quartile) and higher (third quartile) number of expressed molecules. The functional analysis corroborated our hypothesis that miRNA isoforms actively regulate critical genes in cancer. Our results show the activation/deactivation of several critical pathways involved in proliferation, metastasization, tumor immune escape, invasion, and angiogenesis, such as the ILK, HIF1α, and Rac signaling pathways. We then explored the canonical miRNAs/miRNA isoforms’ ability to cluster cancer samples across cohorts, benchmarking three different sets of molecules according to specific RNA modifications. Moving on from using only canonical miRNAs (*CAN* set) to employ all expressed molecules (*ISO* set - canonical miRNAs and miRNA isoforms) allowed us to gain a higher cluster fragmentation that reflected an in-depth clinical-pathological stratification. Notably, in the *ISO_wo_SNV- and ISO*-based clustering results, the TARGET-AML cancer samples were significantly separated into two subclusters characterized by patients without (better prognosis) and with FLT3-ITD mutation (increased relapse risk and reduced overall survival). In the TCGA-COAD and TCGA-READ cohorts, cancer samples were clustered according to lymphatic invasion. Nonetheless, the three sets exclusively clustered cancer samples in TCGA-HNSC (*CAN* set), TCGA-KIRP (*ISO_wo_SNV* set), TCGA-COAD, TCGA-READ, and TCGA-LUSC (*ISO* set). Overall, the combination of canonical miRNAs/miRNA isoforms (*ISO* set) boosted the quality of our results, uniquely outlining clinical-pathological features in cohorts where the other sets failed. Altogether, our results depicted a more complex scenario in which canonical miRNAs and miRNA isoforms seemed to work tightly together to uncover the underlying histopathological differences among cancers. Thus, excluding one of the two may substantially limit our understanding of tumor heterogeneity.

Both canonical miRNAs and miRNA isoforms resulted significantly dysregulated across all cohorts/cancer tissues. These results suggest that miRNA isoforms are not the product of Drosha or Dicer’s arbitrary cleavage, but they are actively expressed and dysregulated across several human cancers. Of the 573 canonical miRNAs dysregulated across cohorts/cancer tissues, we identified 104 characterized by an opposite expression trend compared to their miRNA isoforms. Supported by a previous study (61), we investigated as the first case study the downregulated canonical miR-101-3p and its upregulated shifted isomiR (one nt longer at 5’-end, and two nts shorted at 3’-end) in lung adenocarcinoma primary tumors (TCGA-LUAD). Although it was previously proved that the most robust binding miRNA∷mRNA occurs when the first miRNA nucleotide is U, opposite to an A on mRNA strand (known as t1A) (62), we demonstrated that the canonical miR-101-3p, which starts with U, and the isomiR miR-101-3p (−1|−2), which start with G, are correctly loaded by AGO2 (Figure S5C-D), and are functional molecules with unique targets. Aiming at assessing differences in targeting efficiency, we examined dysregulated and predicted gene targets for the two molecules. We chose PTGS2 (COX-2), an oncogene in lung cancer (39) which is a validated canonical miR-101-3p target gene in different cancers (37,38). Although the predicted binding sites for both miRNA molecules and PTGS2’s 3’ UTR were comparable in terms of binding free energy, we experimentally proved that the sole canonical miR-101-3p targets PTGS2, corroborating the difference in terms of gene targeting for the miRNA molecules. Conversely, DSC2, a protein involved in cell adhesion and often downregulated in cancer (42), is exclusively targeted by miR-101-3p (−1|−2).

In the second case study, we assessed the targetome shifting between two downregulated molecules, the canonical miR-381-3p and its edited form (A-to-I editing site at position 4). In our results, the edited miR-381-3p resulted among the most downregulated molecules across cohorts/cancer tissues. While the role of the canonical miR-381-3p is broadly acknowledged as a tumor suppressor (63), particularly in breast cancer (64), very little is known about the edited form in cancer (25,26). We investigate SYT13 in breast cancer (TCGA-BRCA), an oncogenic gene in different cancers (45,46,65). Unlike the canonical miR-381-3p, which did not show any binding site, our predictions and experiments outlined the ability of the edited miR-381-3p to exclusively regulate the SYT13 expression, suggesting it as a potential tumor suppressor in breast cancer. We experimentally validated an exclusive target (UBE2C) for miR-381-3p to point out the difference in targeting for the two molecules.

Finally, the Overall Survival (OS) and Relapse Free Survival (RFS) results highlighted the importance of including miRNA isoforms over the sole canonical miRNAs. Benchmarking the two sets, we increased the number of significant prognostic signatures from 4 (OS) and 3 (RFS) using canonical miRNA to 9 (OS) and 8 (RFS) employing miRNA isoforms. Interestingly, almost all the potential signatures showed unique molecules, except for the RFS signatures in TCGA-ACC, in which both sets led to a common canonical miRNA, hsa-let-7c-3p.

In conclusion, even though their role is still not well understood, miRNA isoforms may somehow work together with canonical miRNAs to support their function. Through these novel potential diagnostic and prognostic cancer biomarkers, we may be able to shine additional light on those mechanisms related to cancer progression by studying gene regulation via the wider miRNAome.

## ACKNOWLEDGEMENTS

The results published here are in part based upon data generated by The Cancer Genome Atlas (TCGA) Research Network (https://cancer.gov/tcga, DBGap Project ID: 11332) and by the Therapeutically Applicable Research to Generate Effective Treatments initiative (TARGET) of the NCI (http://ocg.cancer.gov/programs/target, DBGap Project ID: 22219). We thank Dr. Andrea Ventura for his valuable comments and suggestions during the development of this study and the scientific discussions during the preparation of the manuscript. We want to thank the Cancer IT Operation Group of The Ohio State University and Mr. Thomas Moore for his valuable technical assistance. We also want to thank the Ohio Supercomputer Center for its resources and technical support. Finally, we thank the Genomics Shared Resource at The Ohio State University Comprehensive Cancer Center (OSU CCC), Columbus, OH, for conducting the qRT-PCR experiments supported in part by the OSU CCC and the National Institutes of Health grant P30 CA16058. M.A. was supported by CTSA award No. UL1TR002649 and NCATS 5KL2TR002648. This work was supported by the National Cancer Institute (National Institute of Health) grant R35CA197706 to C.M.C.

## AUTHOR CONTRIBUTIONS

Conceptualization, R.D and G.N.; Methodology, R.D and G.N.; Software, R.D.; Validation, R.D., L.T., G.L.R.V., P.G. and G.N.; Formal Analysis, R.D. and G.N.; Investigation, R.D., L.T., and G.N.; Resources, R.D., L.T., and G.N.; Data Curation, R.D.; Writing – Original Draft, R.D., L.T., G.L.R.V., and G.N.; Writing – Review & Editing, R.D., L.T., G.L.R.V., P.G., Y.X., M.B., G.P.M., P.F., A.L., M.A., Q.M., G.N., and C.M.C.; Visualization, R.D.; Supervision, G.N. and C.M.C.; Project Administration, C.M.C.; Funding Acquisition, C.M.C.

## REFERENCES

1. Bartel DP. MicroRNAs: genomics, biogenesis, mechanism, and function. Cell. 2004;116:281–97.

2. Mendell JT, Olson EN. MicroRNAs in Stress Signaling and Human Disease. Cell. 2012;148:1172–87.

3. Croce CM. Causes and consequences of microRNA dysregulation in cancer. Nat Rev Genet. 2009;10:704–14.

4. Iorio MV, Croce CM. MicroRNA dysregulation in cancer: diagnostics, monitoring and therapeutics. A comprehensive review. EMBO Mol Med. 2012;4:143–59.

5. Bartel DP. MicroRNAs: Target Recognition and Regulatory Functions. Cell. 2009;136:215–33.

6. Lee LW, Zhang S, Etheridge A, Ma L, Martin D, Galas D, et al. Complexity of the microRNA repertoire revealed by next-generation sequencing. RNA. 2010;16:2170–80.

7. Kozomara A, Griffiths-Jones S. miRBase: annotating high confidence microRNAs using deep sequencing data. Nucl Acids Res. 2014;42:D68–73.

8. Nigita G, Veneziano D, Ferro A. A-to-I RNA Editing: Current Knowledge Sources and Computational Approaches with Special Emphasis on Non-Coding RNA Molecules. Front Bioeng Biotechnol [Internet]. 2015 [cited 2020 Apr 30];3. Available from: http://journal.frontiersin.org/Article/10.3389/fbioe.2015.00037/abstract

9. Nishikura K. A-to-I editing of coding and non-coding RNAs by ADARs. Nat Rev Mol Cell Biol. 2016;17:83–96.

10. Bofill-De Ros X, Kasprzak WK, Bhandari Y, Fan L, Cavanaugh Q, Jiang M, et al. Structural Differences between Pri-miRNA Paralogs Promote Alternative Drosha Cleavage and Expand Target Repertoires. Cell Reports. 2019;26:447–459.e4.

11. Tan GC, Chan E, Molnar A, Sarkar R, Alexieva D, Isa IM, et al. 5′ isomiR variation is of functional and evolutionary importance. Nucleic Acids Res. 2014;42:9424–35.

12. Saunders MA, Liang H, Li W-H. Human polymorphism at microRNAs and microRNA target sites. Proceedings of the National Academy of Sciences. 2007;104:3300–5.

13. Yang W, Chendrimada TP, Wang Q, Higuchi M, Seeburg PH, Shiekhattar R, et al. Modulation of microRNA processing and expression through RNA editing by ADAR deaminases. Nat Struct Mol Biol. 2006;13:13–21.

14. Gallo A, Locatelli F. ADARs: allies or enemies? The importance of A-to-I RNA editing in human disease: from cancer to HIV-1. Biological Reviews. 2012;87:95–110.

15. Maas S, Kawahara Y, Tamburro KM, Nishikura K. A-to-I RNA Editing and Human Disease. RNA Biology. 2006;3:1–9.

16. Dominissini D, Moshitch-Moshkovitz S, Amariglio N, Rechavi G. Adenosine-to-inosine RNA editing meets cancer. Carcinogenesis. 2011;32:1569–77.

17. Kawahara Y, Zinshteyn B, Sethupathy P, Iizasa H, Hatzigeorgiou AG, Nishikura K. Redirection of Silencing Targets by Adenosine-to-Inosine Editing of miRNAs. Science. 2007;315:1137–40.

18. Brennecke J, Stark A, Russell RB, Cohen SM. Principles of MicroRNA–Target Recognition. James C. Carrington, editor. PLoS Biol. 2005;3:e85.

19. Park J-E, Heo I, Tian Y, Simanshu DK, Chang H, Jee D, et al. Dicer recognizes the 5′ end of RNA for efficient and accurate processing. Nature. 2011;475:201–5.

20. Landgraf P, Rusu M, Sheridan R, Sewer A, Iovino N, Aravin A, et al. A Mammalian microRNA Expression Atlas Based on Small RNA Library Sequencing. Cell. 2007;129:1401–14.

21. Wyman SK, Knouf EC, Parkin RK, Fritz BR, Lin DW, Dennis LM, et al. Post-transcriptional generation of miRNA variants by multiple nucleotidyl transferases contributes to miRNA transcriptome complexity. Genome Research. 2011;21:1450–61.

22. Cloonan N, Wani S, Xu Q, Gu J, Lea K, Heater S, et al. MicroRNAs and their isomiRs function cooperatively to target common biological pathways. Genome Biol. 2011;12:R126.

23. Salem O, Erdem N, Jung J, Münstermann E, Wörner A, Wilhelm H, et al. The highly expressed 5’isomiR of hsa-miR-140-3p contributes to the tumor-suppressive effects of miR-140 by reducing breast cancer proliferation and migration. BMC Genomics. 2016;17:566.

24. Kuo W-T, Su M-W, Lee Y, Chen C-H, Wu C-W, Fang W-L, et al. Bioinformatic Interrogation of 5p-arm and 3p-arm Specific miRNA Expression Using TCGA Datasets. JCM. 2015;4:1798–814.

25. Wang Y, Xu X, Yu S, Jeong KJ, Zhou Z, Han L, et al. Systematic characterization of A-to-I RNA editing hotspots in microRNAs across human cancers. Genome Res. 2017;27:1112–25.

26. Pinto Y, Buchumenski I, Levanon EY, Eisenberg E. Human cancer tissues exhibit reduced A-to-I editing of miRNAs coupled with elevated editing of their targets. Nucleic Acids Research. 2018;46:71–82.

27. Loher P, Londin ER, Rigoutsos I. IsomiR expression profiles in human lymphoblastoid cell lines exhibit population and gender dependencies. Oncotarget [Internet]. 2014 [cited 2020 Apr 30];5. Available from: http://www.oncotarget.com/fulltext/2405

28. Telonis AG, Loher P, Jing Y, Londin E, Rigoutsos I. Beyond the one-locus-one-miRNA paradigm: microRNA isoforms enable deeper insights into breast cancer heterogeneity. Nucleic Acids Res. 2015;43:9158–75.

29. Telonis AG, Magee R, Loher P, Chervoneva I, Londin E, Rigoutsos I. Knowledge about the presence or absence of miRNA isoforms (isomiRs) can successfully discriminate amongst 32 TCGA cancer types. Nucleic Acids Research. 2017;45:2973–85.

30. Lu Y, Baras AS, Halushka MK. miRge 2.0 for comprehensive analysis of microRNA sequencing data. BMC Bioinformatics. 2018;19:275.

31. Delaunay J. Prognosis of inv(16)/t(16;16) acute myeloid leukemia (AML): a survey of 110 cases from the French AML Intergroup. Blood. 2003;102:462–9.

32. Rubnitz JE. Favorable Impact of the t(9;11) in Childhood Acute Myeloid Leukemia. Journal of Clinical Oncology. 2002;20:2302–9.

33. Meshinchi S, Woods WG, Stirewalt DL, Sweetser DA, Buckley JD, Tjoa TK, et al. Prevalence and prognostic significance of Flt3 internal tandem duplication in pediatric acute myeloid leukemia. Blood. 2001;97:89–94.

34. Zhang X, He X, Liu Y, Zhang H, Chen H, Guo S, et al. MiR-101-3p inhibits the growth and metastasis of non-small cell lung cancer through blocking PI3K/AKT signal pathway by targeting MALAT-1. Biomedicine & Pharmacotherapy. 2017;93:1065–73.

35. Llorens F, Bañez-Coronel M, Pantano L, del Río JA, Ferrer I, Estivill X, et al. A highly expressed miR-101 isomiR is a functional silencing small RNA. BMC Genomics. 2013;14:104.

36. Distefano R, Nigita G, Veneziano D, Romano G, Croce CM, Acunzo M. isoTar: Consensus Target Prediction with Enrichment Analysis for MicroRNAs Harboring Editing Sites and Other Variations. In: Laganà A, editor. MicroRNA Target Identification [Internet]. New York, NY: Springer New York; 2019. page 211–35. Available from: http://link.springer.com/10.1007/978-1-4939-9207-2_12

37. Chakrabarty A, Tranguch S, Daikoku T, Jensen K, Furneaux H, Dey SK. MicroRNA regulation of cyclooxygenase-2 during embryo implantation. Proceedings of the National Academy of Sciences. 2007;104:15144–9.

38. Hao Y, Gu X, Zhao Y, Greene S, Sha W, Smoot DT, et al. Enforced Expression of miR-101 Inhibits Prostate Cancer Cell Growth by Modulating the COX-2 Pathway *In Vivo*. Cancer Prev Res. 2011;4:1073–83.

39. Liu B, Qu L, Yan S. Cyclooxygenase-2 promotes tumor growth and suppresses tumor immunity. Cancer Cell Int. 2015;15:106.

40. Cui T, Chen Y, Yang L, Mireskandari M, Knösel T, Zhang Q, et al. Diagnostic and prognostic impact of desmocollins in human lung cancer. J Clin Pathol. 2012;65:1100–6.

41. Lin X, Luo W, Wang H, Li R, Huang Y, Chen L, et al. The Role of Prostaglandin-Endoperoxide Synthase-2 in Chemoresistance of Non-Small Cell Lung Cancer. Front Pharmacol. 2019;10:836.

42. Kolegraff K, Nava P, Helms MN, Parkos CA, Nusrat A. Loss of desmocollin-2 confers a tumorigenic phenotype to colonic epithelial cells through activation of Akt/β-catenin signaling. Margolis B, editor. MBoC. 2011;22:1121–34.

43. Xue Y, Xu W, Zhao W, Wang W, Zhang D, Wu P. miR-381 inhibited breast cancer cells proliferation, epithelial-to-mesenchymal transition and metastasis by targeting CXCR4. Biomedicine & Pharmacotherapy. 2017;86:426–33.

44. Mo C, Gao L, Zhu X, Wei K, Zeng J, Chen G, et al. The clinicopathological significance of UBE2C in breast cancer: a study based on immunohistochemistry, microarray and RNA-sequencing data. Cancer Cell Int. 2017;17:83.

45. Kanda M, Shimizu D, Tanaka H, Tanaka C, Kobayashi D, Hayashi M, et al. Synaptotagmin XIII expression and peritoneal metastasis in gastric cancer. British Journal of Surgery. 2018;105:1349–58.

46. Zhang L, Fan B, Zheng Y, Lou Y, Cui Y, Wang K, et al. Identification SYT13 as a novel biomarker in lung adenocarcinoma. J Cell Biochem. 2020;121:963–73.

47. Sun G, Yan J, Noltner K, Feng J, Li H, Sarkis DA, et al. SNPs in human miRNA genes affect biogenesis and function. RNA. 2009;15:1640–51.

48. Cesarini V, Silvestris DA, Tassinari V, Tomaselli S, Alon S, Eisenberg E, et al. ADAR2/miR-589-3p axis controls glioblastoma cell migration/invasion. Nucleic Acids Research. 2018;46:2045–59.

49. Nigita G, Distefano R, Veneziano D, Romano G, Rahman M, Wang K, et al. Tissue and exosomal miRNA editing in Non-Small Cell Lung Cancer. Sci Rep. 2018;8:10222.

50. Shoshan E, Mobley AK, Braeuer RR, Kamiya T, Huang L, Vasquez ME, et al. Reduced adenosine-to-inosine miR-455-5p editing promotes melanoma growth and metastasis. Nat Cell Biol. 2015;17:311–21.

51. Guo L, Li Y, Cirillo KM, Marick RA, Su Z, Yin X, et al. mi-IsoNet: systems-scale microRNA landscape reveals rampant isoform-mediated gain of target interaction diversity and signaling specificity. Briefings in Bioinformatics. 2021;bbab091.

52. Loher P, Karathanasis N, Londin E, Bray P, Pliatsika V, Telonis AG, et al. IsoMiRmap—fast, deterministic, and exhaustive mining of isomiRs from short RNA-seq datasets. Gorodkin J, editor. Bioinformatics. 2021;btab016.

53. Bizuayehu TT, Lanes CF, Furmanek T, Karlsen BO, Fernandes JM, Johansen SD, et al. Differential expression patterns of conserved miRNAs and isomiRs during Atlantic halibut development. BMC Genomics. 2012;13:11.

54. Neilsen CT, Goodall GJ, Bracken CP. IsomiRs –the overlooked repertoire in the dynamic microRNAome. Trends in Genetics. 2012;28:544–9.

55. Katoh T, Hojo H, Suzuki T. Destabilization of microRNAs in human cells by 3′ deadenylation mediated by PARN and CUGBP1. Nucleic Acids Res. 2015;43:7521–34.

56. Bail S, Swerdel M, Liu H, Jiao X, Goff LA, Hart RP, et al. Differential regulation of microRNA stability. RNA. 2010;16:1032–9.

57. Brodersen P, Voinnet O. Revisiting the principles of microRNA target recognition and mode of action. Nat Rev Mol Cell Biol. 2009;10:141–8.

58. Moore MJ, Scheel TKH, Luna JM, Park CY, Fak JJ, Nishiuchi E, et al. miRNA–target chimeras reveal miRNA 3′-end pairing as a major determinant of Argonaute target specificity. Nat Commun. 2015;6:8864.

59. Van der Kwast RVCT, Woudenberg T, Quax PHA, Nossent AY. MicroRNA-411 and Its 5′-IsomiR Have Distinct Targets and Functions and Are Differentially Regulated in the Vasculature under Ischemia. Molecular Therapy. 2020;28:157–70.

60. Katoh T, Sakaguchi Y, Miyauchi K, Suzuki T, Kashiwabara S -i., Baba T, et al. Selective stabilization of mammalian microRNAs by 3’ adenylation mediated by the cytoplasmic poly(A) polymerase GLD-2. Genes & Development. 2009;23:433–8.

61. Li Z, Qu Z, Wang Y, Qin M, Zhang H. miR-101-3p sensitizes non-small cell lung cancer cells to irradiation. Open Medicine. 2020;15:413–23.

62. Gebert LFR, MacRae IJ. Regulation of microRNA function in animals. Nat Rev Mol Cell Biol. 2019;20:21–37.

63. Qiao G, Li J, Wang J, Wang Z, Bian W. miR‑381 functions as a tumor suppressor by targeting ETS1 in pancreatic cancer. Int J Mol Med [Internet]. 2019 [cited 2021 Mar 2]; Available from: http://www.spandidos-publications.com/10.3892/ijmm.2019.4206

64. Yi D, Xu L, Wang R, Lu X, Sang J. miR-381 overcomes cisplatin resistance in breast cancer by targeting MDR1: miR-381 overcomes cisplatin resistance. Cell Biol Int. 2019;43:12–21.

65. Li Z, Zhu W, Xiong L, Yu X, Chen X, Lin Q. Role of high expression levels of STK39 in the growth, migration and invasion of non-small cell type lung cancer cells. Oncotarget. 2016;7:61366–77.

