## Supplementary Information for "A concurrent canonical and modified miRNAome pan-cancer study on TCGA and TARGET cohorts leads to an enhanced resolution in cancer"

#### List of Supplementary Material


1. **Supplementary Methods**
2. **Supplementary Figures**
    - Figure S1. Distribution of expressed molecules across cohorts and RNA modification types, Related to Figure 1
    - Figure S2. Significant Pathways Enriched in at Least Five Cohorts/Cancer Tissues, Related to Figure 1
    - Figure S3. Sample Visualization and Clustering Workflow, Related to Figure 2
    - Figure S4. Dataset-based Clustering Comparison, Related to Figure 2
    - Figure S5. Transfections, RNA-binding Protein Immunoprecipitation (RIP), and Binding Sites, Related to Figure 4
    - Figure S6. Risk Score-Based Prognostic Signature Discovery Workflow, Related to Figure 5
3. **Supplementary Tables**
    - Table S1. Primers list for cloning and sequencing of target genes 3' UTRs
    - Table S2. A-to-I RNA editing sites
    - Table S3. Expressed canonical miRNAs/miRNA isoforms distribution over modification types and miRNA arms

- Table S4. Enriched Pathways Across Cohorts/Cancer Tissues
- Table S5. Clustering clinical-pathological features
- Table S6. Dysregulated miRNA isoforms across cohort/cancer tissues
- Table S7. Distribution of dysregulated miRNA isoform across modification types
- Table S8. Dysregulated miRNA isoforms with opposite trends
- Table S9. Dysregulated genes supplied with predicted targets for the selected case studies
- Table S10. Risk score-based signatures list

**4. Supplementary References**

### 1. Supplementary Methods

#### Data and Code Availability

The data and source code produced in this work will be stored on Code Ocean and Zenodo repositories. *The Cancer Genome Atlas* (TCGA) (1,2) and *The Therapeutically Applicable Research to Generate Effective Treatments* (TARGET) (3) miRNA-Seq samples, patients' clinical-pathological data, and somatic mutations used during this study can be found at http://portal.gdc.cancer.gov. Moreover, additional single nucleotide DNA variants were downloaded from COSMIC (4) at https://cancer.sanger.ac.uk/cosmic and dbSNP (5) at https://ftp.ncbi.nih.gov/snp/archive/b154/VCF, while A-to-I miRNA Editing sites were downloaded from MiREDiBase (6).

#### MiRNA-Seq Data

The TCGA (v20) and TARGET (v20) miRNA-Seq samples (BAM file format) were retrieved via the Genomic Data Commons Data Portal (GDC Data Portal, https://gdc-portal.nci.nih.gov). Following the authorization from the data access committee (DBGap Project IDs: 11332 and 22219 for TCGA and TARGET repositories, respectively), samples were downloaded checking the following options: "*sequencing reads*" (Data Category), "*miRNA-Seq*" (Experimental Strategy), "*bam*" (Data Format), and "*TCGA*" and "*TARGET*" (Program). We adopted the TCGA and TARGET barcode to pair samples and clinical data, processing 33 TCGA (10,977 samples over 10,250 adult patients) and 5 TARGET (2,373 samples over 1,123 pediatric patients) cohorts. Both datasets provide insights into a wide range of adult and pediatric cancers and are considered the most prominent and reliable public resources.

#### Clinical Data

The GDC Data Portal provides cohort patients' clinical data in *JavaScript Object Notation* (JSON) file format (e.g., *clinical.cart.XXXX.json*). Clinical data files are available within the GDC *cart* section along with the miRNA-Seq samples. In addition, the GDC Legacy Archive (https://portal.gdc.cancer.gov/legacy-archive) offers additional TARGET clinical data in Microsoft Excel file format. Collected clinical data files were downloaded, opportunely parsed, and harmonized across cohorts.

**Single Nucleotide DNA/RNA Variant Data**

After creating the proper login account, COSMIC data (v92) were downloaded by clicking on "*Data*" and "*Downloads*" (main menu), and then on the "*CosmicNCV.tsv.gz*" link located on the right side ("*Non coding variants*" section). The dbSNP (v154) database (NCBI SNP), a VCF file format, was downloaded via the following link: *https://ftp.ncbi.nih.gov/snp/archive/b154/VCF/GCF_000001405.38.gz*. Finally, A-to-I miRNA editing sites were downloaded from MiREDiBase (6).

**MiRNA-Seq Quality Check (QC)**

Downloaded BAM files were converted into FASTQ files by leveraging the *bamToFastq* tool (*bedtools* v2.25.0 package) (7) and finally quality-filtered through the ConDeTri tool (v2.3) (8) (parameters: -pb=fq -lq=20 -hq=30 -minlen=15 -sc=33). The workflow is summarized in Figure 1A.

**MiRNA Isoforms Mapping and Quantification**

All quality-filtered sequences were aligned to the Ensembl (www.ensembl.org) human genome (hg38) and annotated via an in-house designed workflow (Figure 1A). The workflow itself leveraged the miRge 2.0 (9), a pipeline for canonical miRNAs/miRNA isoforms annotation based on the latest miRBase (v22) (10) and miRGeneDB 2.0 (11) datasets. The miRge 2.0 pipeline represents one of the major pipelines for canonical miRNAs/miRNA isoforms profiling (12–14), given its reliability in identifying A-to-I RNA editing sites and accuracy comparable to a well-established miRNA editing detection approach (15).

We performed miRge 2.0 using the default parameters. The annotation process covered a wide range of molecules (Figure 1B). The annotation process covered both shifted and non-shifted isomiRs. The last ones were characterized by sequence shifting affecting the 5', 3', or both ends (addition/trimming of nucleotides with respect to their canonical miRNA sequence). The process also annotated molecules subjected to Single Nucleotide Variants (i.e., SNPs, A-to-I RNA Editing sites, somatic mutations), hereafter called SNVs. At first, the workflow leveraged the miRge 2.0 pipeline. It then extended the annotation results with additional information, such as the latest known SNPs (dbSNP), somatic mutations (COSMIC, TCGA, and TARGET), and A-to-I RNA Editing sites collected from 40 studies (Table S2), to maximize accuracy. Finally, a data filtering phase discarded all those annotated miRNA isoforms having SNVs not yet characterized (unknown) involving the first or last two nucleotides (Figure 1A).

Finally, data were collected into tab-separated text files per cohort (i.e., TCGA/Lung Adenocarcinoma shortened TCGA-LUAD).

#### MiRNA Isoform Nomenclature

We designed a unique human-readable way to label each mapped miRNA isoform in this work. A label is *de facto* a combination of the canonical miRNA, pre-miRNA, 5'- and 3'-end shifting, and a "*Compact Idiosyncratic Gapped Alignment Report*" string, shortened CIGAR (https://samtools.github.io/hts-specs/SAMv1.pdf). Succinctly, the CIGAR string is a standard to indicate base match/mismatch and other operations (e.g., insertion - *I* - and deletion - *D*), used to describe sequence alignments. However, it is essential to note that miRge current implementation supports only *insertion* operations, potentially posing limitations on which miRNA isoforms can or cannot be mapped.

To better understand how a label looks like, let us consider the *miR-21-5p__mir-21__-1__+1__2MG21M* label. It represents a mapped miR-21-5p isoform (pre-miRNA: mir-21), which undergoes the following modifications:

- A single genomic nucleotide shifting (left) at 5'-end, denoted by *-1*;
- A single genomic nucleotide shifting (right) at 3'-end, denoted by *+1*;
- A single nucleotide modification at position 3 (i.e., an A-to-I RNA editing site), represented using the CIGAR string *2MG21M*. The *2M* and *21M* indicate a perfect match between the miRNA isoform and the genomic reference, while at position 3 (after 2M, or two matches), we ended up with a G instead of A (reference).

Above all, we extended the miRge CIGAR string paradigm, replacing each insertion (denoted with *I*), which solely involves the miRNA isoform extremities, with the mapped read's corresponding nucleotide. We adopted such an extension to deal with those miRNA isoforms (same miRNA, pre-miRNA, and shifting) which show identical CIGAR string but undergo different single nucleotide insertions (not reported by miRge). For the sake of clarity, we report below an example of two miRNA isoforms with a single nucleotide insertion at 5'-end and no shifting at 3'-end (represented by *0*), which undergo a single nucleotide insertion at 5'-end:

- **miRge CIGAR**:
    - MiRNA isoform n°1:
        - Canonical miRNA: miR-30a-5p
        - Pre-miRNA: mir-30a
        - Sequence mapped: ATGTAAACATCCTCGACTGGAAG
        - CIGAR: I22M
        - 5'-end shifting: -1
        - 3'-end shifting: 0

        - Tag: miR-30a-5p__ mir-30a__-1__0__ I22M
    - MiRNA isoform n°2:
        - Canonical miRNA: miR-30a-5p
        - Pre-miRNA: mir-30a
        - Sequence mapped: TTGTAAACATCCTCGACTGGAAG
        - CIGAR: I22M
        - 5'-end shifting: -1
        - 3'-end shifting: 0
        - Tag: miR-30a-5p__ mir-30a__-1__0__ I22M
- **miRge CIGAR extension**:
    - MiRNA isoform n°1:
        - Canonical miRNA: miR-30a-5p
        - Pre-miRNA: mir-30a
        - Sequence mapped: ATGTAAACATCCTCGACTGGAAG
        - CIGAR: A22M
        - 5'-end shifting: -1
        - 3'-end shifting: 0
        - Tag: miR-30a-5p__ mir-30a__-1__0__ A22M
    - MiRNA isoform n°2:
        - Canonical miRNA: miR-30a-5p
        - Pre-miRNA: mir-30a
        - Sequence mapped: TTGTAAACATCCTCGACTGGAAG
        - CIGAR: T22M
        - 5'-end shifting: -1

- 3’-end shifting: 0
- Tag: miR-30a-5p__ mir-30a__-1__0__ T22M

Given the example above, it is natural and straightforward to link the label *miR-21-5p__mir-21__0__0__22M* to the canonical miRNA form. Furthermore, the way we label mapped miRNA isoforms should give users a clear hint on which modifications occur, along with their precise location within the sequence. Finally, all novel miRNA molecules were labeled as follows: *miR-n##*, where ### represents a positive incremental integer (e.g., miR-n86-3p and miR-n136-5p).

**Data Aggregation, Annotation, and Filtering**

For each dataset/cohort, multiple samples per patient and tissue type (i.e., a patient with more than one Primary Solid Tumor sample) were aggregated by calculating the raw read counts expression average. Then, raw read counts tables were enriched with additional information, including known SNVs, such as SNPs from dbSNP, somatic mutations from COSMIC, TCGA, and TARGET, along with 2,885 A-to-I miRNA Editing sites. A final filter discarded miRNA isoforms with not yet characterized (unknown) SNVs involving the first or last two nucleotides. We removed these molecules because these unknown SNVs could be due to potential sequencing errors or imperfections in the linker ligation during the construction of the cDNA library (16–18). The workflow is summarized in Figure 1A.

**Data Normalization**

In this study, every downstream analysis was preceded by a normalization process. In this process, raw read counts were filtered and normalized, applying the described criterion. Depending on the type of data to be normalized, we used *Reads Per Million miRNA mapped reads (RPM)* for miRNA isoforms and *Fragments Per Kilobase Million (FPKM)* for transcripts. Specifically for miRNA isoforms, we computed the RPM expression from the raw read counts. An expression filtering was then applied to retain miRNA isoforms/genes having a minimum expression of $\lfloor(\text{RPM}|\text{FPKM})\ \text{geometric mean}\rfloor > 1$. Lastly, the set of expression-filtered molecules was used to extract the corresponding raw read counts from the initial table and finally normalized via the *trimmed mean of M-values* (TMM) method (19) using the *calcNormFactors* function available in *edgeR* (v3.24.3) (20,21), an R (v3.4.4) *BioConductor* (v3.6) (22) package.

**High Dimensionality Reduction and Clustering**

Aimed to investigate the benefits and drawbacks of using specific miRNA isoforms for clustering purposes, we benchmarked three sets of expressed molecules (minimum expression of $\lfloor\text{RPM geometric mean}\rfloor > 1$) grouped according to their modification type (Figure S3A). In the first set, labeled "*CAN*," we considered only canonical miRNAs (miRBase v22). In the second one, marked "*ISO_wo_SNV*," we used both canonical miRNAs and shifted isomiRs with no SNVs. In the last set, labeled "*ISO*," we considered all expressed canonical miRNAs and isomiRs, including the shifted ones. We applied an in-house designed workflow (Figure S3B) to each set, aiming to assess molecules' ability to cluster samples across different cohorts/cancer tissues.

The workflow starts extracting all expressed miRNA isoforms from each cancer cohort (e.g., cancer samples in TCGA-LUAD) according to a minimum expression of $\lfloor\text{RPM geometric mean}\rfloor > 1$. It then collects the raw reads count expression of the extracted miRNA isoforms, collapsing the data into a single massive raw read counts table and normalizing it via TMM (expressed molecules as rows, cohorts' cancer samples as columns).

Afterward, a nonlinear dimensionality-reduction technique (Uniform Manifold Approximation and Projection - UMAP) (23) is applied to reduce high-dimensional data (the normalized table) into two-dimensional data, leveraging the Bray-Curtis distance (24). Except for the distance (metric=braycurtis),

we performed UMAP (*umap-learn* Python package, v0.4.6) using its default parameters (min_dist=.1, n_neighbors=15, n_components=2), specifying a seed (random_state=99) for reproducibility purposes. The resulting lowered-dimension table is used for sample visualization and clustering analysis (see Figure S3). Next, we applied the DBSCAN algorithm (25) to perform an unsupervised clustering using the Euclidean distance and the UMAP two-dimensional data as input. In a nutshell, the DBSCAN algorithm (*scikit-learn* Python package, v0.22.1) mainly relies on two distinct parameters: *eps* and *min_samples*. The *eps* value represents the maximum distance between two points, used to decide whether to group them or not. The *min_samples* value represents the minimum number of nearest neighbors (samples) for a point to be considered as a core point. Due to DBSCAN's high sensibility to the *eps* value, the workflow calculates the optimal value using the *K-Nearest Neighbor Distance*. Having optimal *eps* speeds up the analysis, allowing us to explore only the *min_samples* parameter (from 3 to 25, with incremental steps of 1). All clustering results were evaluated via Adjusted Rand Index (ARI) (26), Adjusted Mutual Information (AMI) (27), and Fowlkes-Mallows Index (FMI) (28) scores.
Finally, all results were filtered considering a percentage of mislabeled samples (noise) less than 5%.

**Differentially miRNA Isoforms Expression Analysis**

To detect dysregulated abundant miRNA isoforms across cancer cohorts, we performed a differential expression (DE) analysis with a minimum of 5 samples per cohort/tissue type: i.e., TCGA-LUAD/Solid Normal Tissue and TCGA-LUAD/Primary Solid Tumor. Input data were normalized according to the normalization criterion described in the "*Data Normalization*" section. Linear *fold change* and statistical significance were calculated using the mean and the two-sided unpaired Mann-Whitney U test (29). The resulting p-values were adjusted using Benjamini-Hochberg's correction (30), using the *fdrcorrection* function from the *statsmodels* (v0.11.1) Python package. Finally, dysregulated molecules with *adjusted p-value* < 0.05 and |*linear fold change*| > 1.5 were retained.

**Differentially Genes Expression Analysis**

Unlike the previous section, we performed the gene differential expression analysis for the pathways enrichment analysis and each molecule in the two case studies: the canonical miR-101-3p, its shifted isomiR, and the canonical miR-381-3p and its edited form.

We grouped samples according to the first (Q1) and third (Q3) quartile of each molecule case study, investigating dysregulated genes to assess potential target variability. Instead, in the pathway enrichment analysis, we grouped each cohort's cancer samples into two groups: low (Q1) and high (Q3) abundant expressed miRNA isoforms.

Input data were normalized according to the normalization criterion described in the "*Data Normalization*" section. Then, the gene differential expression analysis was performed using *edgeR* (v3.32.1), a *Bioconductor* (v3.6) R (v3.4.4) package, keeping all those dysregulated genes with *adjusted p-value* <0.05 and |*linear fold change*| >1.5.

**Pathway Enrichment Analysis**

The analysis relied on differentially expressed genes generated as described above. We then employed the Ingenuity® Pathway Analysis (IPA) software (v01-16) to perform a pathway enrichment analysis for each cohort/cancer tissue. Finally, we retained all those pathways characterized by $|\text{z-score}| \geq 2$ and *p-value* <0.01. We generated a heatmap plotting the z-scores of the most significant pathways enriched in at least five cohorts/cancer tissues, clustering them via the Bray-Curtis distance metric and Complete Linkage method. The heatmap was created using the *clustermap* function from the *seaborn* (v0.10.1) Python package.

**Risk Score-based Prognostic Signature**

We individually reviewed the clinical data associated with each cohort, considering the patient's survival time independent from each other and treating the censoring time as right-censored data. We included only those patients with survival times greater than zero.

We designed a 2-stages workflow (Figure S6) for optimal prognostic signature estimation to assess miRNA isoforms' effectiveness as prognostic biomarkers. The workflow relies on 10-fold cross-validation (CV) strategy to avoid potential bias and model under-/over-fitting.

The first stage arranges the patients' miRNA isoforms expression data. It starts by checking whether the cohort/cancer tissue provides at least *twenty* patients per event type (e.g., *dead* and *alive* for the Overall Survival), representing a minimum number for reliable results. Then, the RPM expression data is extracted for the selected patients. Finally, the stage normalizes via TMM the raw read counts expression of the retained expressed miRNA isoforms and calculates a correlation matrix from the normalized data.

The second stage explores potential prognostic signatures by initially removing highly correlated miRNA isoforms. The stage iteratively retains those molecules with a correlation coefficient lower than a specific *threshold*, from 0.5 to 0.8, with incremental steps of 0.01. A recursive feature elimination and cross-validation selection of the best number of features (RFECV) is then applied to rank miRNA isoforms, using Logistic Regression as the estimator (GridSearchCV for optimal logistic regression parameters estimation), from the *scikit-learn* Python package (v0.22.1).

On this reduced set of data, the 2-phases 10-fold CV takes place. Besides the correlation coefficient threshold and RFECV, each CV iteration implements the same set of steps. We employed a stratified shuffle splitting policy to ensure a proportional balance of event/nonevent patients among the training (66% of patients) and validation (34% of patients) sets.

The first phase focuses on extracting a significant prognostic signature (*P-value* <0.05) from the training set and uses it for validation. It generates a univariate Cox proportional hazard regression model to assess the relationship between miRNA isoforms expression and patients' survival. Observed *Cox p-values* are then adjusted using Benjamini-Hochberg's *correction*. It next performs a multivariate Cox proportional hazard regression model based on the most significant (*Cox FDR* <0.05) miRNA isoforms extracted from the univariate model. Both univariate and multivariate Cox regression models leverage the *CoxPHFitter* function from the *lifelines* (v0.25.4) Python package. Extracted miRNA isoforms are narrowed down by keeping those molecules with *Cox FDR* <0.05 (multivariate model) and further reduced by applying a feature selection strategy. The selection strategy relies on RFECV, coupled with a Logistic Regression Model used as the estimator.

Next, a *Risk Score* is computed for each patient by linearly combining the expression value and the regression coefficient (univariate model) related to the reduced set of miRNA isoforms (selection strategy), as shown below:

$$Risk\ Score = \sum_{i=1}^{n} exp_i \cdot \beta_i$$

Where *n* represents the number of reduced miRNA isoforms, *exp* the expression, and *ß* the regression coefficient (univariate Cox model). Patients are then separated into high- and low-risk groups using the patients' Risk Score median as a cutoff. Finally, the last step ends by generating a Kaplan-Meier Plot (31), a p-value (Log Rank Test), and a prognostic signature accuracy (area under the curve - AUC) based on the two risk groups. For the Kaplan-Meier Plot, we leveraged the *survfit* and *ggsurvplot* functions provided by the *survival* (v3.2-3) R (v3.4.4) package.

The second phase checks the training signature significance (*P-value* <0.05), starting a new CV iteration if the significance is not achieved. The second phase follows a similar set of steps as the previous one, focusing only on those miRNA isoforms identified by the training signature. At the end of the second phase, the signature significance is evaluated (*P-value* <0.05) for the validation set, and if significant, the signature is temporarily held. After completing the ten CV iterations, the workflow extracts all miRNA isoforms occurring in at least 50% of held signatures. The signature-based on the most frequent miRNA isoforms is finally tested on the entire dataset (training + validation sets). Significance and AUC values are stored, and the second stage is reiterated on the next correlation coefficient *threshold*.

The workflow concludes by picking up the prognostic signature with the highest AUC for each cohort/cancer tissue. In this analysis, we investigate both *Overall Survival* (*OS*) (event: *death*, nonevent: *alive*) and *Relapse Free Survival* (RFS) (event: *relapse*, nonevent: *no relapse*).

**MiRNA Isoform-Target Prediction**

All miRNA isoform-target predictions were generated by isoTar v1.2.1 (Distefano et al., 2019). IsoTar leverages state-of-the-art prediction tools, such as miRmap (v1.1) (33), TargetScan (v7.0) (34), PITA (v6) (35), RNAhybrid (v2.1.2) (36), and miRanda (v3.3a) (37). IsoTar focuses solely on seed regions of 7-8 nucleotides in length (7mer-A1, 7mer-m8, 8mer), with no mismatch or G:U base pairs (wobbles). All predictions were performed using isoTar default parameters.

#### 2. Supplementary Figures

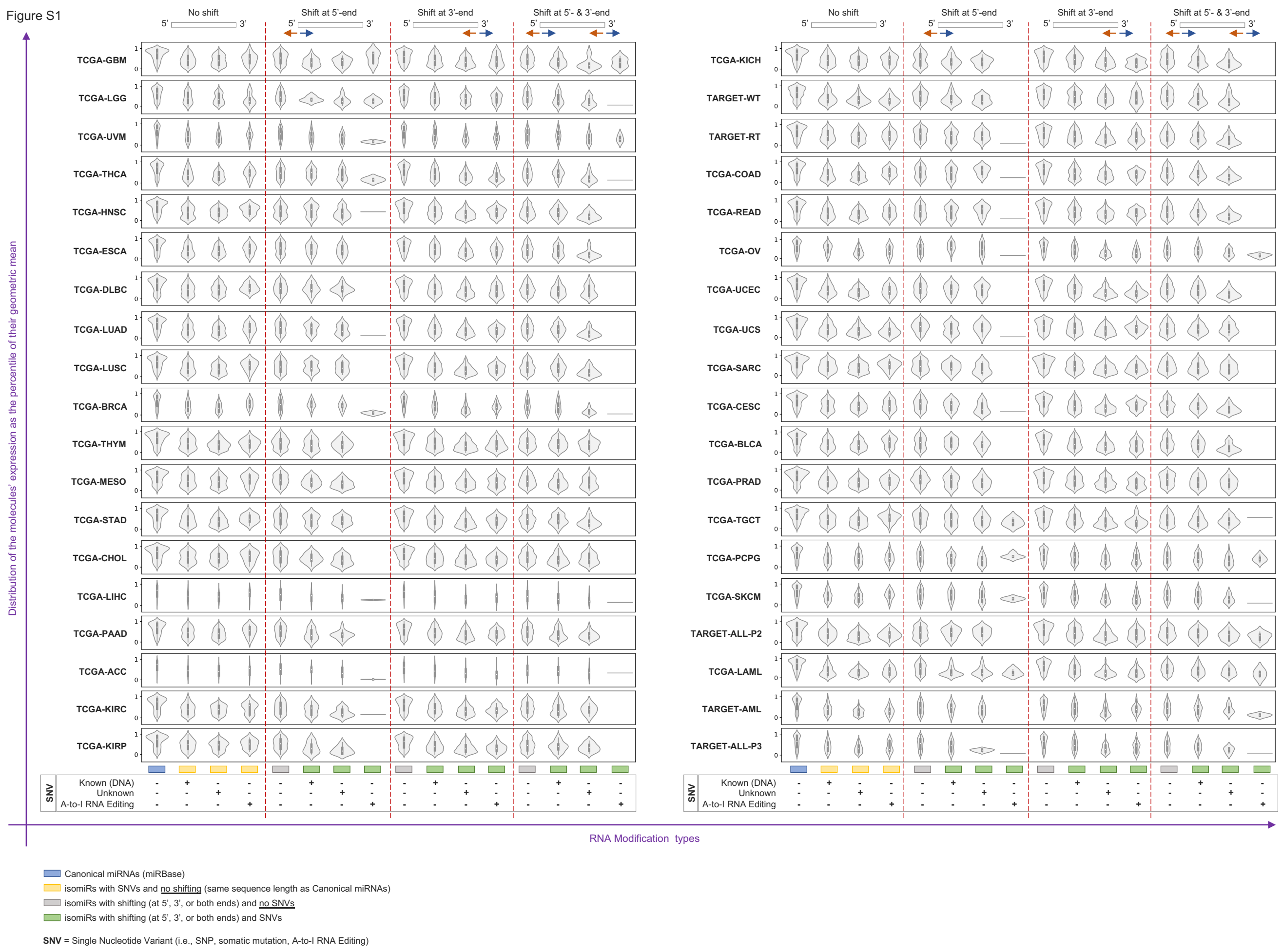


**Figure S1. Distribution of expressed molecules across cohorts and RNA modification types, Related to Figure 1**

The figure shows the distribution of the molecules' expression as the percentile of their geometric mean across cohorts and RNA modification types.

Figure S2

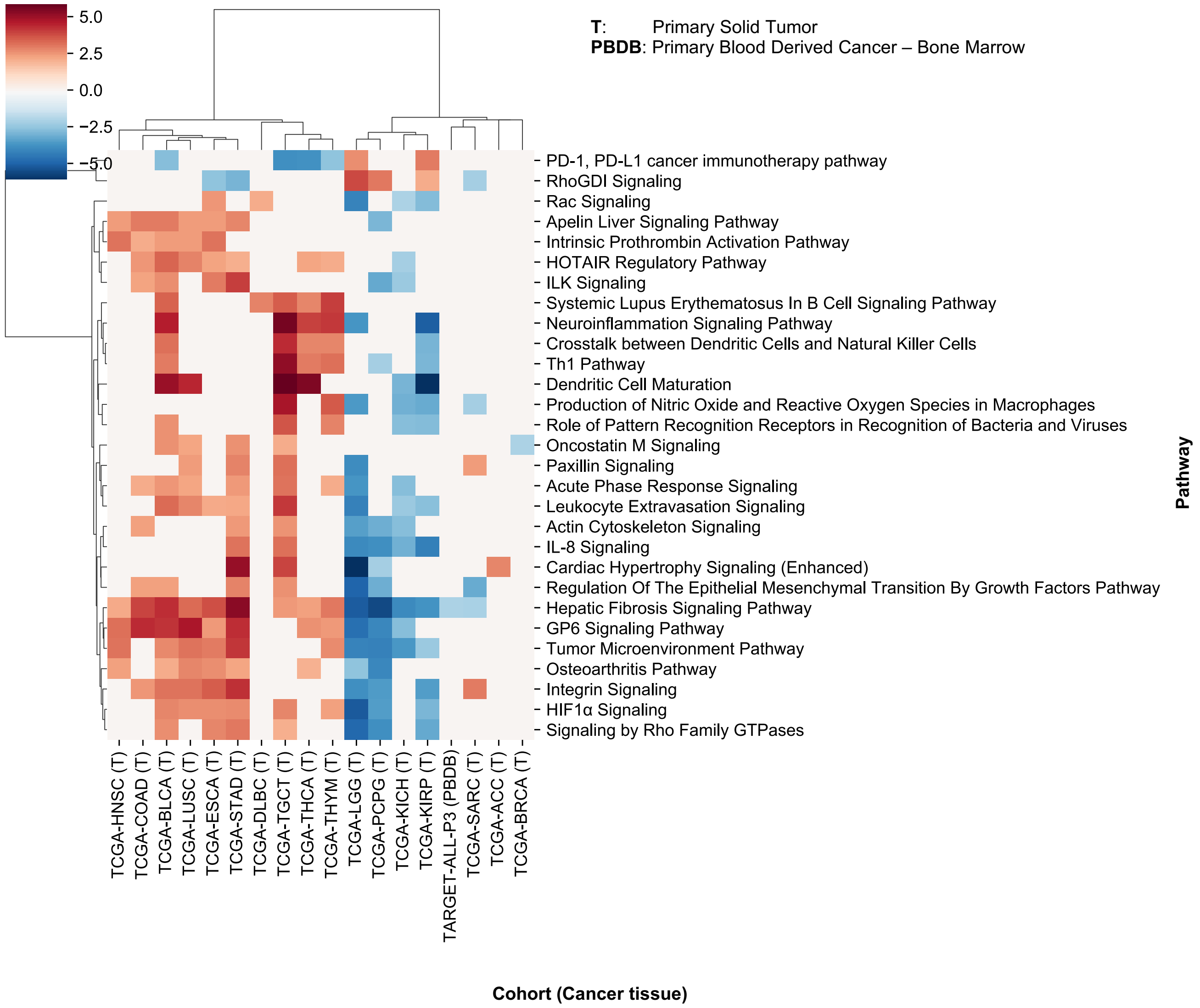
5.0
2.5
0.0
−2.5
−5.0
T: Primary Solid Tumor
PBDB: Primary Blood Derived Cancer – Bone Marrow
PD-1, PD-L1 cancer immunotherapy pathway
RhoGDI Signaling
Rac Signaling
Apelin Liver Signaling Pathway
Intrinsic Prothrombin Activation Pathway
HOTAIR Regulatory Pathway
ILK Signaling
Systemic Lupus Erythematosus In B Cell Signaling Pathway
Neuroinflammation Signaling Pathway
Crosstalk between Dendritic Cells and Natural Killer Cells
Th1 Pathway
Dendritic Cell Maturation
Production of Nitric Oxide and Reactive Oxygen Species in Macrophages
Role of Pattern Recognition Receptors in Recognition of Bacteria and Viruses
Oncostatin M Signaling
Paxillin Signaling
Acute Phase Response Signaling
Leukocyte Extravasation Signaling
Actin Cytoskeleton Signaling
IL-8 Signaling
Cardiac Hypertrophy Signaling (Enhanced)
Regulation Of The Epithelial Mesenchymal Transition By Growth Factors Pathway
Hepatic Fibrosis Signaling Pathway
GP6 Signaling Pathway
Tumor Microenvironment Pathway
Osteoarthritis Pathway
Integrin Signaling
HIF1α Signaling
Signaling by Rho Family GTPases
Pathway
TCGA-HNSC (T)
TCGA-COAD (T)
TCGA-BLCA (T)
TCGA-LUSC (T)
TCGA-ESCA (T)
TCGA-STAD (T)
TCGA-DLBC (T)
TCGA-TGCT (T)
TCGA-THCA (T)
TCGA-THYM (T)
TCGA-LGG (T)
TCGA-PCPG (T)
TCGA-KICH (T)
TCGA-KIRP (T)
TARGET-ALL-P3 (PBDB)
TCGA-SARC (T)
TCGA-ACC (T)
TCGA-BRCA (T)
Cohort (Cancer tissue)

**Figure S2. Significant Pathways Enriched in at Least Five Cohorts/Cancer Tissues, Related to Figure 1**

The figure shows significant ($|z\text{-}score| \geq 2$ and *p-value* <0.01) pathways enriched in at least five cohorts/cancer tissues. Pathways and cohorts/cancer tissues are clustered according to the Bray-Curtis distance metric and Complete Linkage method. See Supplementary Information for more details.

Figure S3

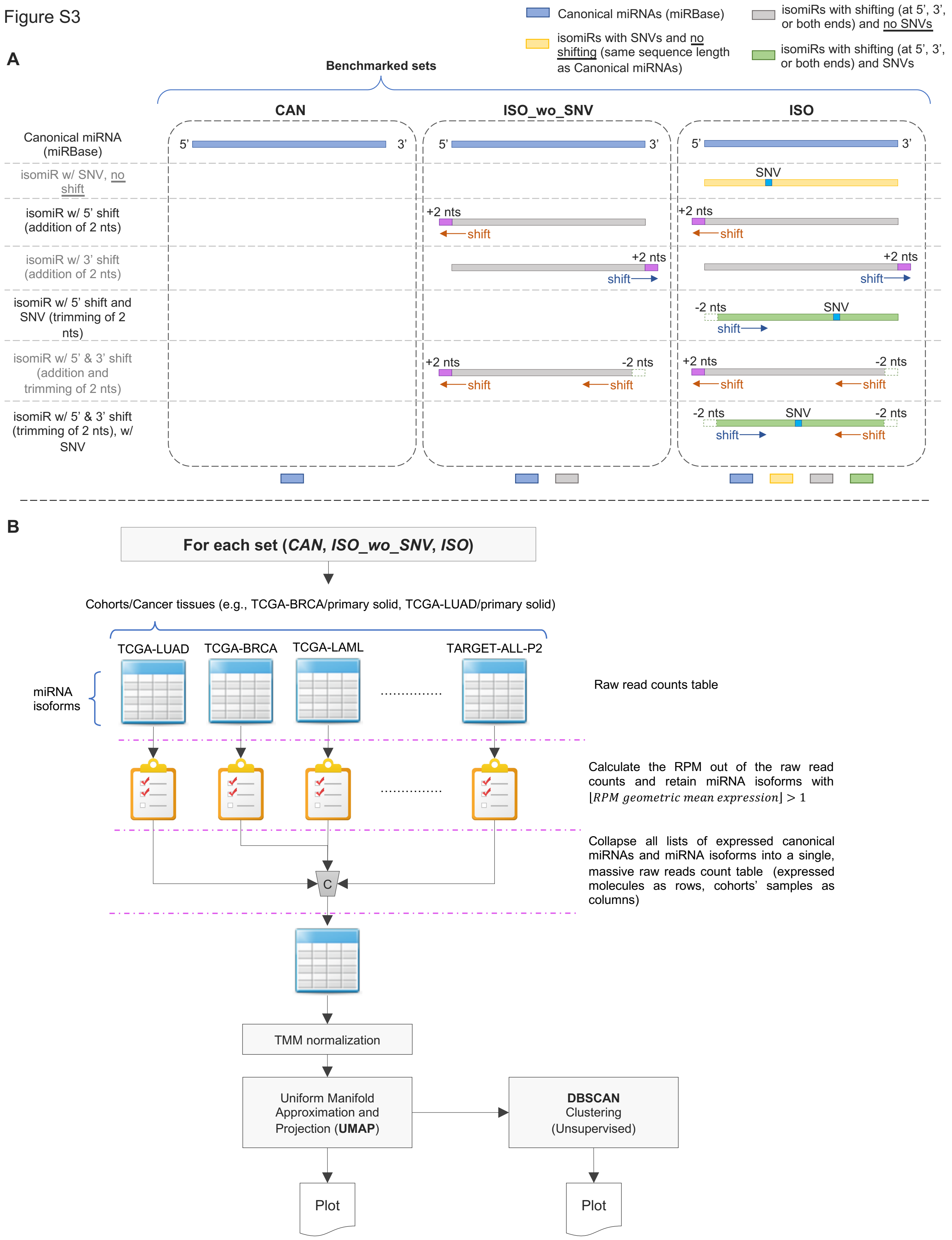

Canonical miRNAs (miRBase)
isomiRs with SNVs and no shifting (same sequence length as Canonical miRNAs)
isomiRs with shifting (at 5', 3', or both ends) and no SNVs
isomiRs with shifting (at 5', 3', or both ends) and SNVs
A
Benchmarked sets
CAN
ISO_wo_SNV
ISO
Canonical miRNA (miRBase)
isomiR w/ SNV, no shift
isomiR w/ 5' shift (addition of 2 nts)
isomiR w/ 3' shift (addition of 2 nts)
isomiR w/ 5' shift and SNV (trimming of 2 nts)
isomiR w/ 5' & 3' shift (addition and trimming of 2 nts)
isomiR w/ 5' & 3' shift (trimming of 2 nts), w/ SNV
5'
3'
SNV
+2 nts
-2 nts
shift
B
For each set (*CAN*, *ISO_wo_SNV*, *ISO*)
Cohorts/Cancer tissues (e.g., TCGA-BRCA/primary solid, TCGA-LUAD/primary solid)
TCGA-LUAD
TCGA-BRCA
TCGA-LAML
TARGET-ALL-P2
miRNA isoforms
Raw read counts table
Calculate the RPM out of the raw read counts and retain miRNA isoforms with $\lfloor RPM\ geometric\ mean\ expression \rfloor > 1$
Collapse all lists of expressed canonical miRNAs and miRNA isoforms into a single, massive raw reads count table (expressed molecules as rows, cohorts' samples as columns)
C
TMM normalization
Uniform Manifold Approximation and Projection (**UMAP**)
**DBSCAN** Clustering (Unsupervised)
Plot
Plot

**Figure S3. Sample Visualization and Clustering Workflow, Related to Figure 2**

(A-B) The figure shows how we defined the three sets to benchmark (A) and the designed workflow used for visualization and benchmarking (B). The three sets (*CAN*, *ISO_wo_SNV*, and *ISO*) encompassed expressed molecules based on specific modification types. In particular, the "*CAN*" set considered only canonical miRNAs (miRBase v22); the "*ISO_wo_SNV*" one used both canonical miRNAs and shifted isomiRs without SNVs; the "*ISO*" set employed all expressed canonical miRNAs and isomiRs, including the shifted ones. The workflow extracts expressed molecules from every cohort, condensing the information into a single massive table (expressed molecules as rows, cohorts' samples as columns). A nonlinear dimensionality-reduction technique (Uniform Manifold Approximation and Projection - UMAP) is applied to reduce high-dimensional data into a two-dimensional matrix for data visualization and evaluation. The reduced two-dimensional matrix is then used to perform an unsupervised clustering by leveraging the DBSCAN algorithm to test the sets' clustering capability. See Supplementary Information for more details.

Figure S4

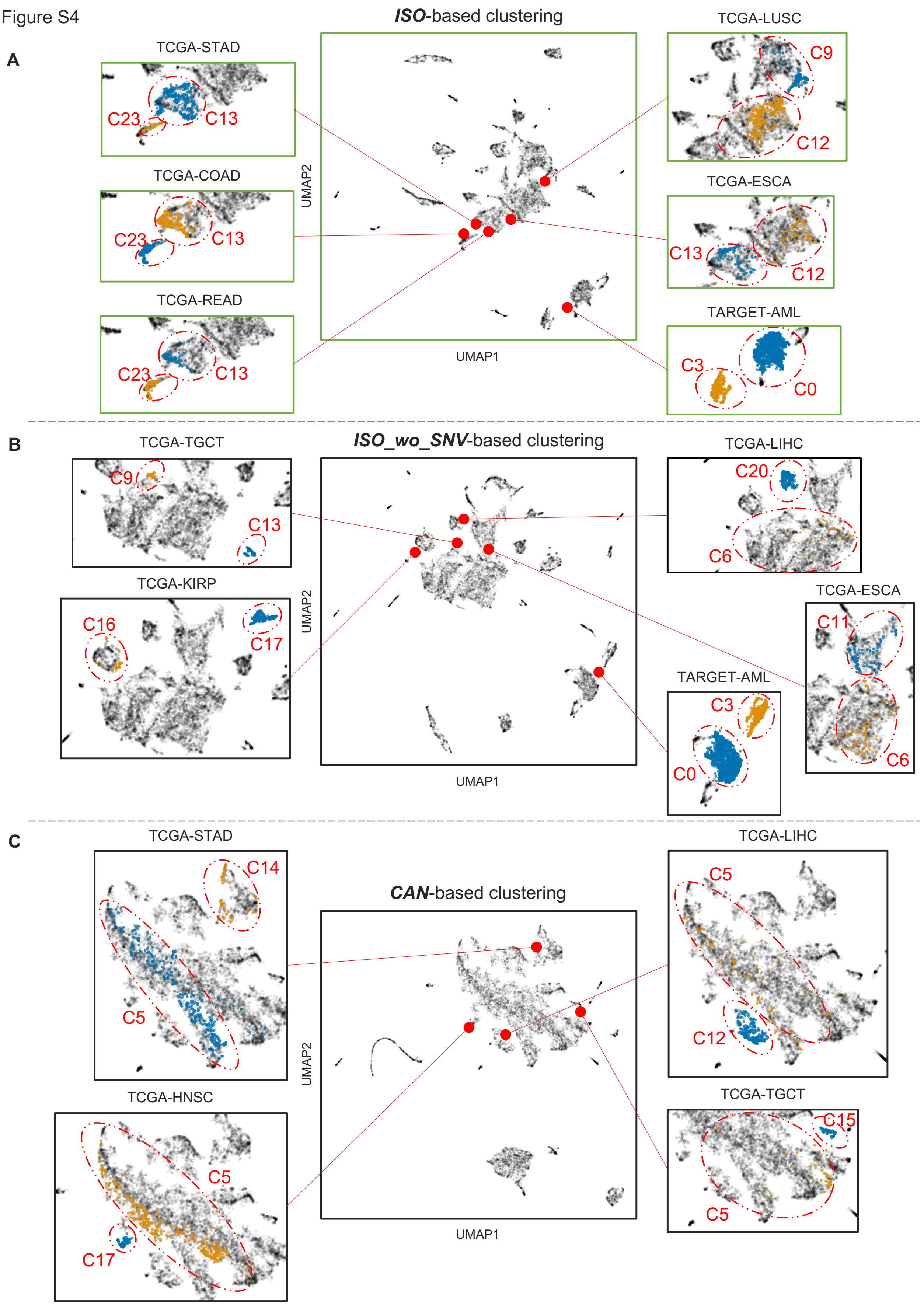

A
ISO-based clustering
TCGA-STAD
C23
C13
TCGA-COAD
C23
C13
TCGA-READ
C23
C13
TCGA-LUSC
C9
C12
TCGA-ESCA
C13
C12
TARGET-AML
C3
C0
UMAP2
UMAP1
B
ISO_wo_SNV-based clustering
TCGA-TGCT
C9
C13
TCGA-KIRP
C16
C17
TCGA-LIHC
C20
C6
TCGA-ESCA
C11
C6
TARGET-AML
C3
C0
UMAP2
UMAP1
C
CAN-based clustering
TCGA-STAD
C14
C5
TCGA-HNSC
C5
C17
TCGA-LIHC
C5
C12
TCGA-TGCT
C15
C5
UMAP2
UMAP1

**Figure S4. Dataset-based Clustering Comparison, Related to Figure 2**

(A-C) Comparison between *ISO*- (A), *ISO_wo_SNV*- (B), and *CAN*-based cancer samples clustering (C). Panels (A-C) display only clusters related to the most prominent and significant clinical-pathological features we considered (Table S5). Each panel shows common and unique cohorts identified by the three sets. Clusters and their IDs are highlighted throughout the figure. Coloring is used to highlight clusters within the same set/cohort.

Figure S5

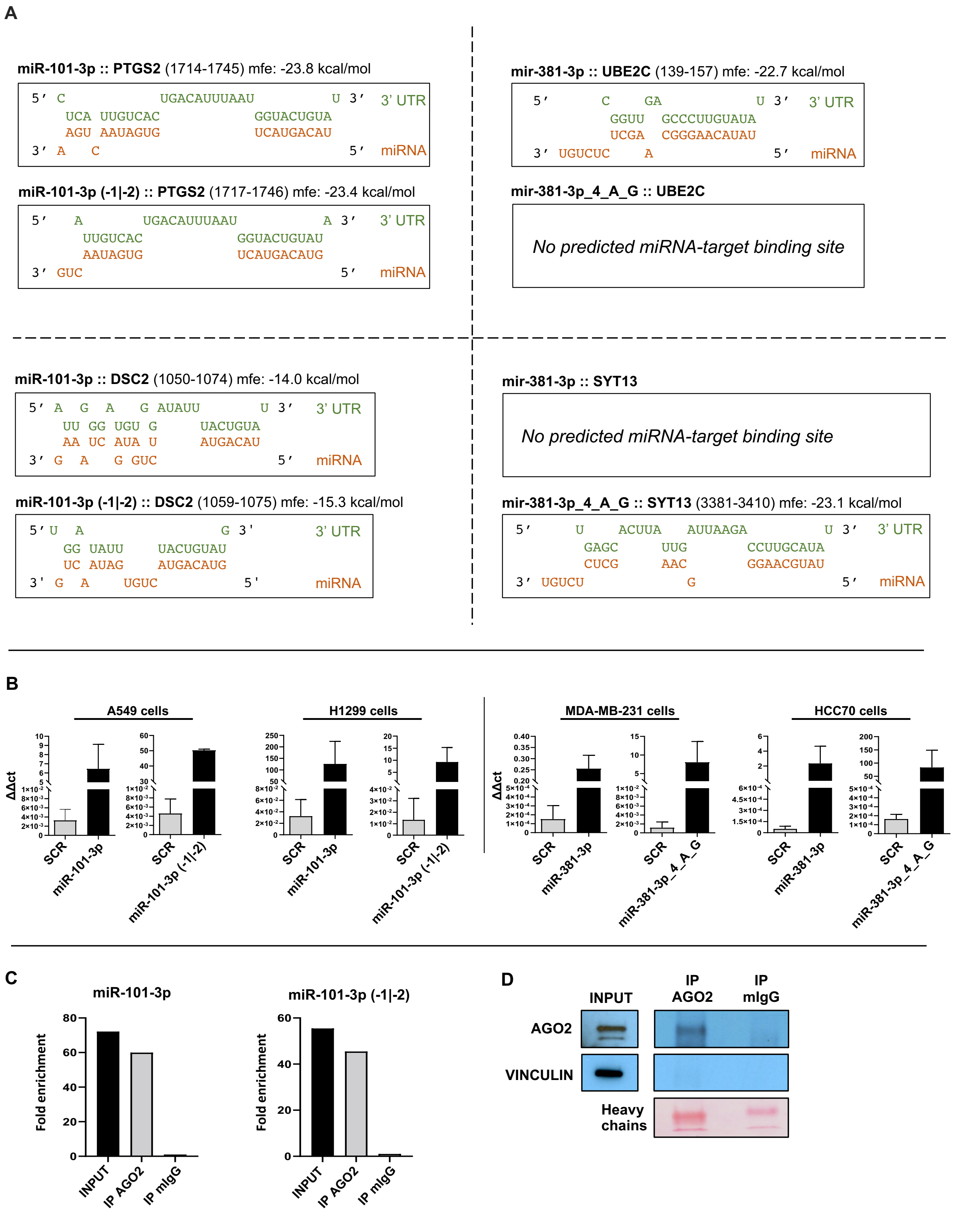
A
miR-101-3p :: PTGS2 (1714-1745) mfe: -23.8 kcal/mol
5' C UGACAUUUAAU U 3' 3' UTR
UCA UUGUCAC GGUACUGUA
AGU AAUAGUG UCAUGACAU
3' A C 5' miRNA
miR-101-3p (-1|-2) :: PTGS2 (1717-1746) mfe: -23.4 kcal/mol
5' A UGACAUUUAAU A 3' 3' UTR
UUGUCAC GGUACUGUAU
AAUAGUG UCAUGACAUG
3' GUC 5' miRNA
mir-381-3p :: UBE2C (139-157) mfe: -22.7 kcal/mol
5' C GA U 3' 3' UTR
GGUU GCCCUUGUAUA
UCGA CGGGAACAUAU
3' UGUCUC A 5' miRNA
mir-381-3p_4_A_G :: UBE2C
No predicted miRNA-target binding site
miR-101-3p :: DSC2 (1050-1074) mfe: -14.0 kcal/mol
5' A G A G AUAUU U 3' 3' UTR
UU GG UGU G UACUGUA
AA UC AUA U AUGACAU
3' G A G GUC 5' miRNA
miR-101-3p (-1|-2) :: DSC2 (1059-1075) mfe: -15.3 kcal/mol
5' U A G 3' 3' UTR
GG UAUU UACUGUAU
UC AUAG AUGACAUG
3' G A UGUC 5' miRNA
mir-381-3p :: SYT13
No predicted miRNA-target binding site
mir-381-3p_4_A_G :: SYT13 (3381-3410) mfe: -23.1 kcal/mol
5' U ACUUA AUUAAGA U 3' 3' UTR
GAGC UUG CCUUGCAUA
CUCG AAC GGAACGUAU
3' UGUCU G 5' miRNA
B
A549 cells
H1299 cells
MDA-MB-231 cells
HCC70 cells
ΔΔct
SCR
miR-101-3p
miR-101-3p (-1|-2)
miR-381-3p
miR-381-3p_4_A_G
C
miR-101-3p
miR-101-3p (-1|-2)
Fold enrichment
INPUT
IP AGO2
IP mIgG
D
INPUT
IP AGO2
IP mIgG
AGO2
VINCULIN
Heavy chains

**Figure S5. Transfections, RNA-binding Protein Immunoprecipitation (RIP), and Binding Sites, Related to Figure 4**

(A) Histograms report the expression of miR-101-3p and miR-101-3p (-1|-2) in A549 and H1299 lung cancer cell lines, along with miR-381-3p and miR-381-3p_4_A_G in MDA-MB-231 and HCC70 breast cancer cell lines, after 48 hours from the transfection with specific mirVana™ miRNA mimics for miRNA isoforms and the negative scramble miRNA control.

(B) RIP was performed using A549 cell lysate and either anti-AGO2 or Normal Mouse IgG (negative control) as the immunoprecipitating antibody. Purified RNA was then analyzed by qRT-PCR using Taqman probes specific for miR-101-3p (canonical miRNA) and miR-101-3p (-1|-2). INPUT sample represents total lysate before immunoprecipitation (positive control). Fold enrichment was calculated considering IP IgG as the reference sample.

(C) Western blotting related to the AGO2 protein expression in A549 total lysate (INPUT) and A549 lysates immunoprecipitated with anti-AGO2 and Normal Mouse IgG, respectively.

(D) Graphical representation of the binding site between the four gene targets (PTGS2, DSC2, UBE2C, and SYT13) and the miRNA isoforms of interest.

Figure S6

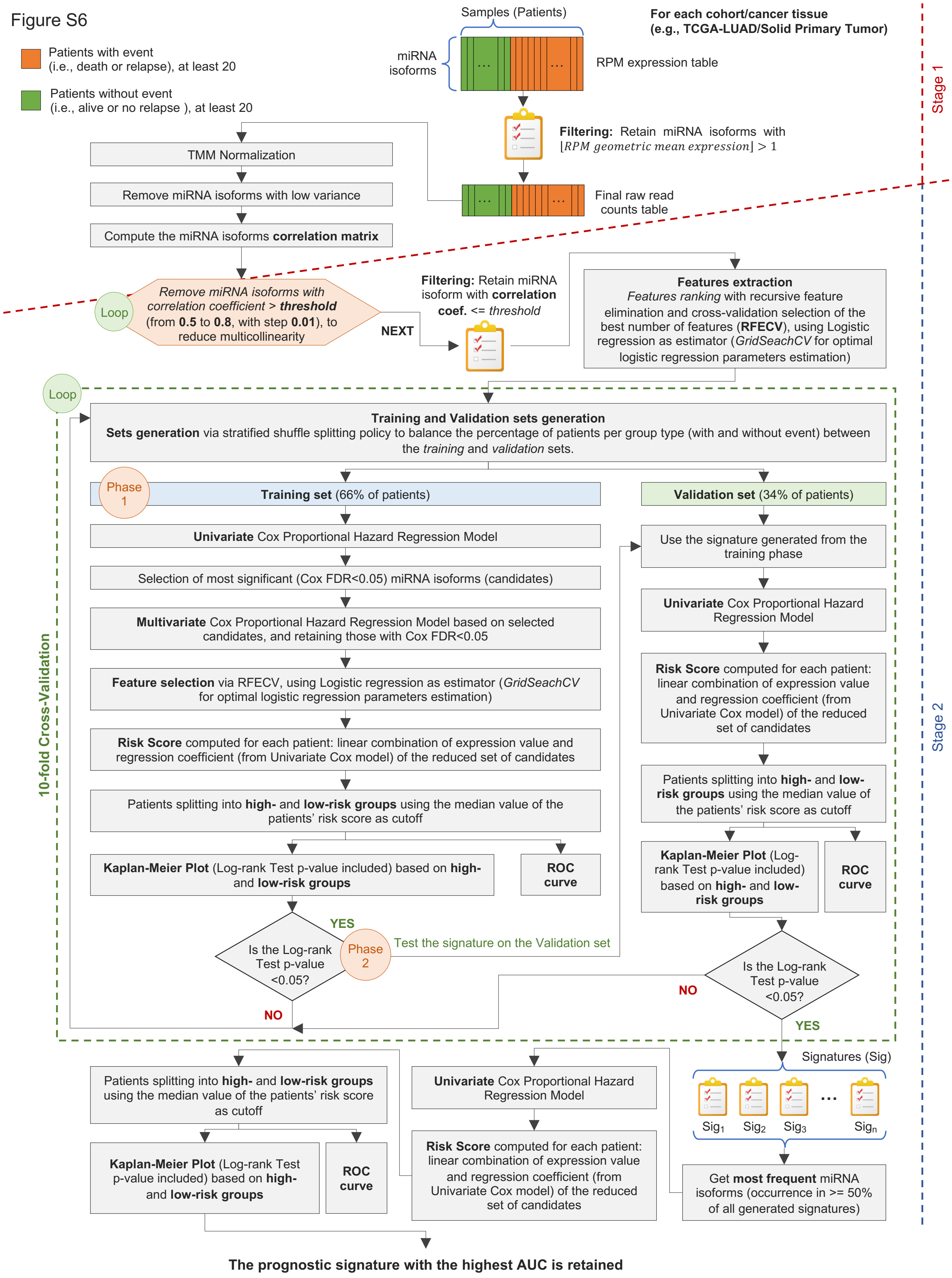
Patients with event
(i.e., death or relapse), at least 20
Patients without event
(i.e., alive or no relapse ), at least 20
Samples (Patients)
miRNA isoforms
RPM expression table
For each cohort/cancer tissue
(e.g., TCGA-LUAD/Solid Primary Tumor)
Stage 1
Filtering: Retain miRNA isoforms with
⌊RPM geometric mean expression⌋ > 1
Final raw read counts table
TMM Normalization
Remove miRNA isoforms with low variance
Compute the miRNA isoforms correlation matrix
Loop
Remove miRNA isoforms with correlation coefficient > threshold (from 0.5 to 0.8, with step 0.01), to reduce multicollinearity
NEXT
Filtering: Retain miRNA isoform with correlation coef. <= threshold
Features extraction
Features ranking with recursive feature elimination and cross-validation selection of the best number of features (RFECV), using Logistic regression as estimator (GridSeachCV for optimal logistic regression parameters estimation)
Loop
10-fold Cross-Validation
Training and Validation sets generation
Sets generation via stratified shuffle splitting policy to balance the percentage of patients per group type (with and without event) between the training and validation sets.
Phase 1
Training set (66% of patients)
Validation set (34% of patients)
Univariate Cox Proportional Hazard Regression Model
Selection of most significant (Cox FDR<0.05) miRNA isoforms (candidates)
Multivariate Cox Proportional Hazard Regression Model based on selected candidates, and retaining those with Cox FDR<0.05
Feature selection via RFECV, using Logistic regression as estimator (GridSeachCV for optimal logistic regression parameters estimation)
Risk Score computed for each patient: linear combination of expression value and regression coefficient (from Univariate Cox model) of the reduced set of candidates
Patients splitting into high- and low-risk groups using the median value of the patients' risk score as cutoff
Kaplan-Meier Plot (Log-rank Test p-value included) based on high- and low-risk groups
ROC curve
Is the Log-rank Test p-value <0.05?
YES
NO
Phase 2
Test the signature on the Validation set
Use the signature generated from the training phase
Univariate Cox Proportional Hazard Regression Model
Risk Score computed for each patient: linear combination of expression value and regression coefficient (from Univariate Cox model) of the reduced set of candidates
Patients splitting into high- and low-risk groups using the median value of the patients' risk score as cutoff
Kaplan-Meier Plot (Log-rank Test p-value included) based on high- and low-risk groups
ROC curve
Is the Log-rank Test p-value <0.05?
NO
YES
Stage 2
Signatures (Sig)
Sig1 Sig2 Sig3 … Sign
Get most frequent miRNA isoforms (occurrence in >= 50% of all generated signatures)
Univariate Cox Proportional Hazard Regression Model
Risk Score computed for each patient: linear combination of expression value and regression coefficient (from Univariate Cox model) of the reduced set of candidates
Patients splitting into high- and low-risk groups using the median value of the patients' risk score as cutoff
Kaplan-Meier Plot (Log-rank Test p-value included) based on high- and low-risk groups
ROC curve
The prognostic signature with the highest AUC is retained

**Figure S6. Risk Score-Based Prognostic Signature Discovery Workflow, Related to Figure 5**

The figure shows the 2-stages workflow designed for prognostic signature discovery for Overall Survival (OS) and Relapse Free Survival (RFS). See Supplementary Information for more details.

#### 3. Supplementary Tables

**Table S1. Primers list for cloning and sequencing of target genes 3' UTRs**

List of primers used for cloning and sequencing experiments. See Supplementary Information for more details.

**Table S2. A-to-I RNA editing sites**

The table reports the list of known A-to-I miRNA Editing sites available in MiREDiBase (v1).

**Table S3. Expressed canonical miRNAs/miRNA isoforms distribution over modification types and miRNA arms**

The table shows the distribution of modification types of expressed molecules over 5p and 3p arms, the shifting extent at 5'- and 3'-ends, along with the number of molecules subjected to SNPs, somatic mutations, and A-to-I RNA editing sites.

**Table S4. Enriched Pathways Across Cohorts/Cancer Tissues**

The table reports significant pathways enriched in at least one cohort/cancer tissue, retained according to $|z\text{-}score| \geq 2$ and *p-value* $<0.01$.

**Table S5. Clustering clinical-pathological features**

The table reports clinical-pathological features per cohort, we the most prominent and significant ones ($Chi\text{-}Square\ p\text{-}value < 0.01$) considered to investigate clustering results from a clinical standpoint. Features are grouped according to each benchmarked set of molecules (*CAN*, *ISO_wo_SNV*, and *ISO*).

**Table S6. Dysregulated miRNA isoforms across cohort/cancer tissues**

The table reports the complete list of dysregulated molecules across cohorts/cancer tissues, retained according to a |*linear fold change*| >1.5 and an *FDR adjusted p-value* <0.05.

**Table S7. Distribution of dysregulated miRNA isoform across modification types**

The table outlines the distribution of dysregulated molecules and modification types across 5p and 3p arms, filtered according to a |*linear fold change*| >1.5 and an *FDR adjusted p-value* <0.05. In addition, the table provides additional information on the distribution of dysregulated molecules with 5'- and 3'-end shifting, SNPs/somatic mutations, and A-to-I RNA editing sites.

**Table S8. Dysregulated miRNA isoforms with opposite trends**

The table reports canonical miRNAs characterized by an opposite expression trend than their miRNA isoforms across cohorts/comparisons. Molecules are retained according to a |*linear fold change*| >1.5 and an *FDR adjusted p-value* <0.05.

**Table S9. Dysregulated genes supplied with predicted targets for the selected case studies**

The table shows the dysregulated genes between the first (Q1) and third (Q3) quartiles of each case study: canonical miRNAs *miR-101-3p* and *miR-381-3p*, isomiR miR-101-3p (-1|-2), and edited miRNA miR-381-3p_4_A_G. Whether available, each dysregulated gene is supplied with a predicted target consensus generated via isoTar, based on five prediction tools: PITA, RNAhybrid, TargetScan, miRanda, and miRmap.

**Table S10. Risk score-based signatures list**

The table reports the most prominent and significant risk score-based signatures for Overall Survival (OS) and Relapse Free Survival (RFS), each one supplied with the molecules list, the area under the curve score (AUC), and p-value. Canonical miRNAs are highlighted in grey.

#### 4. Supplementary References


1. Cancer Genome Atlas Research Network, Weinstein JN, Collisson EA, Mills GB, Shaw KRM, Ozenberger BA, et al. The Cancer Genome Atlas Pan-Cancer analysis project. Nat Genet. 2013;45:1113–20.

2. Tomczak K, Czerwińska P, Wiznerowicz M. Review The Cancer Genome Atlas (TCGA): an immeasurable source of knowledge. wo. 2015;1A:68–77.

3. Gadd S, Huff V, Walz AL, Ooms AHAG, Armstrong AE, Gerhard DS, et al. A Children's Oncology Group and TARGET initiative exploring the genetic landscape of Wilms tumor. Nat Genet. 2017;49:1487–94.

4. Tate JG, Bamford S, Jubb HC, Sondka Z, Beare DM, Bindal N, et al. COSMIC: the Catalogue Of Somatic Mutations In Cancer. Nucleic Acids Research. 2019;47:D941–7.

5. Sherry ST. dbSNP: the NCBI database of genetic variation. Nucleic Acids Research. 2001;29:308–11.

6. Marceca GP, Distefano R, Tomasello L, Lagana' A, Russo F, Calore F, et al. MiREDiBase: a manually curated database of editing events in microRNAs [Internet]. Bioinformatics; 2020 Sep. Available from: http://biorxiv.org/lookup/doi/10.1101/2020.09.04.283689

7. Quinlan AR, Hall IM. BEDTools: a flexible suite of utilities for comparing genomic features. Bioinformatics. 2010;26:841–2.

8. Smeds L, Künstner A. ConDeTri - A Content Dependent Read Trimmer for Illumina Data. Donlin MJ, editor. PLoS ONE. 2011;6:e26314.

9. Lu Y, Baras AS, Halushka MK. miRge 2.0 for comprehensive analysis of microRNA sequencing data. BMC Bioinformatics. 2018;19:275.

10. Kozomara A, Griffiths-Jones S. miRBase: annotating high confidence microRNAs using deep sequencing data. Nucl Acids Res. 2014;42:D68–73.

11. Fromm B, Domanska D, Høye E, Ovchinnikov V, Kang W, Aparicio-Puerta E, et al. MirGeneDB 2.0: the metazoan microRNA complement. Nucleic Acids Research. 2020;48:D132–41.

12. Chu A, Robertson G, Brooks D, Mungall AJ, Birol I, Coope R, et al. Large-scale profiling of microRNAs for The Cancer Genome Atlas. Nucleic Acids Res. 2016;44:e3–e3.

13. Kanke M, Baran-Gale J, Villanueva J, Sethupathy P. miRquant 2.0: an Expanded Tool for Accurate Annotation and Quantification of MicroRNAs and their isomiRs from Small RNA-Sequencing Data. Journal of Integrative Bioinformatics [Internet]. 2016 [cited 2021 Apr 14];13. Available from: https://www.degruyter.com/document/doi/10.1515/jib-2016-307/html

14. Loher P, Karathanasis N, Londin E, Bray P, Pliatsika V, Telonis AG, et al. IsoMiRmap—fast, deterministic, and exhaustive mining of isomiRs from short RNA-seq datasets. Gorodkin J, editor. Bioinformatics. 2021;btab016.

15. Alon S, Erew M, Eisenberg E. DREAM: a webserver for the identification of editing sites in mature miRNAs using deep sequencing data. Bioinformatics. 2015;31:2568–70.

16. de Hoon MJL, Taft RJ, Hashimoto T, Kanamori-Katayama M, Kawaji H, Kawano M, et al. Cross-mapping and the identification of editing sites in mature microRNAs in high-throughput sequencing libraries. Genome Research. 2010;20:257–64.

17. Jones MR, Quinton LJ, Blahna MT, Neilson JR, Fu S, Ivanov AR, et al. Zcchc11-dependent uridylation of microRNA directs cytokine expression. Nat Cell Biol. 2009;11:1157–63.

18. Katoh T, Sakaguchi Y, Miyauchi K, Suzuki T, Kashiwabara S -i., Baba T, et al. Selective stabilization of mammalian microRNAs by 3' adenylation mediated by the cytoplasmic poly(A) polymerase GLD-2. Genes & Development. 2009;23:433–8.

19. Robinson MD, Oshlack A. A scaling normalization method for differential expression analysis of RNA-seq data. Genome Biol. 2010;11:R25.

20. McCarthy DJ, Chen Y, Smyth GK. Differential expression analysis of multifactor RNA-Seq experiments with respect to biological variation. Nucleic Acids Research. 2012;40:4288–97.

21. Robinson MD, McCarthy DJ, Smyth GK. edgeR: a Bioconductor package for differential expression analysis of digital gene expression data. Bioinformatics. 2010;26:139–40.

22. Gentleman RC, Carey VJ, Bates DM, Bolstad B, Dettling M, Dudoit S, et al. Bioconductor: open software development for computational biology and bioinformatics. Genome Biol. 2004;5:R80.

23. McInnes L, Healy J, Melville J. UMAP: Uniform Manifold Approximation and Projection for Dimension Reduction. arXiv:180203426 [cs, stat] [Internet]. 2018 [cited 2020 May 3]; Available from: http://arxiv.org/abs/1802.03426

24. Bray JR, Curtis JT. An Ordination of the Upland Forest Communities of Southern Wisconsin. Ecological Monographs. 1957;27:325–49.

25. Ester M, Kriegel H-P, Sander J, Xu X. A density-based algorithm for discovering clusters in large spatial databases with noise. 1996. page 226–31.

26. Rand WM. Objective Criteria for the Evaluation of Clustering Methods. Journal of the American Statistical Association. 1971;66:846–50.

27. Vinh NX, Epps J, Bailey J. Information theoretic measures for clusterings comparison: is a correction for chance necessary? Proceedings of the 26th Annual International Conference on Machine Learning - ICML '09 [Internet]. Montreal, Quebec, Canada: ACM Press; 2009 [cited 2020 Dec 29]. page 1–8. Available from: http://portal.acm.org/citation.cfm?doid=1553374.1553511

28. Fowlkes EB, Mallows CL. A Method for Comparing Two Hierarchical Clusterings. Journal of the American Statistical Association. 1983;78:553–69.

29. Mann HB, Whitney DR. On a Test of Whether one of Two Random Variables is Stochastically Larger than the Other. Ann Math Statist. 1947;18:50–60.

30. Benjamini Y, Hochberg Y. Controlling the False Discovery Rate: A Practical and Powerful Approach to Multiple Testing. Journal of the Royal Statistical Society: Series B (Methodological). 1995;57:289–300.

31. Kaplan EL, Meier P. Nonparametric Estimation from Incomplete Observations. Journal of the American Statistical Association. 1958;53:457–81.

32. Distefano R, Nigita G, Veneziano D, Romano G, Croce CM, Acunzo M. isoTar: Consensus Target Prediction with Enrichment Analysis for MicroRNAs Harboring Editing Sites and Other Variations. In: Laganà A, editor. MicroRNA Target Identification [Internet]. New York, NY: Springer New York; 2019 [cited 2021 Mar 1]. page 211–35. Available from: http://link.springer.com/10.1007/978-1-4939-9207-2_12

33. Vejnar CE, Zdobnov EM. miRmap: Comprehensive prediction of microRNA target repression strength. Nucleic Acids Research. 2012;40:11673–83.

34. Lewis BP, Burge CB, Bartel DP. Conserved Seed Pairing, Often Flanked by Adenosines, Indicates that Thousands of Human Genes are MicroRNA Targets. Cell. 2005;120:15–20.

35. Kertesz M, Iovino N, Unnerstall U, Gaul U, Segal E. The role of site accessibility in microRNA target recognition. Nat Genet. 2007;39:1278–84.

36. Heyne S, Costa F, Rose D, Backofen R. GraphClust: alignment-free structural clustering of local RNA secondary structures. Bioinformatics. 2012;28:i224–32.

37. John B, Enright AJ, Aravin A, Tuschl T, Sander C, Marks DS. Human MicroRNA Targets. James C. Carrington, editor. PLoS Biol. 2004;2:e363.
